# Comparative genomics in plant fungal pathogens (Mycosphaerellaceae): variation in mitochondrial composition due to at least five independent intron invasions

**DOI:** 10.1101/694562

**Authors:** Juliana E. Arcila, Rafael E. Arango, Javier M. Torres, Tatiana Arias

## Abstract

Fungi provide new opportunities to study highly differentiated mitochondrial DNA. Mycosphaerellaceae is a highly diverse fungal family containing a variety of pathogens affecting many economically important crops.

Mitochondria plays a major role in fungal metabolism and fungicide resistance but up until now only two annotated mitochondrial genomes have been published in this family. We sequenced and annotated mitochondrial genomes of selected Mycosphaerellaceae species that diverged ∼66 MYA. During this time frame, mitochondrial genomes expanded significantly due to at least five independent invasions of introns into different electron transport chain genes. Comparative analysis revealed high variability in size and gene order among mitochondrial genomes even of closely related organisms, truncated extra gene copies and, accessory genes in some species. Gene order variability was common probably due to rearrangements caused by mobile intron invasion. Three three *cox1* copies and bicistronic transcription of *nad2-nad3* and *atp6-atp8* in *Pseudocercospora fijiensis* were confirmed experimentally. Even though we found variation in mitochondrial genome composition, there was no evidence of hybridization when comparing nuclear and mitochondrial dataset sets for fungal plant pathogens analyzed here. Disentangling the causes of variation in mitochondrial genome composition in plant pathogenic fungal move us closer to understanding the molecular mechanisms responsible for vital functions in fungi ultimately aiding in controlling these diseases.

## 1. Introduction

From their origin as an early alpha proteobacterial endosymbiont to their current state as cellular organelles, large scale genomic reorganization has taken place in the mitochondria of all main eukaryotic lineages (Aguileta *et al*., 2014). Although fungi are a lineage more closely related to animals (Clark-Walker, 1992; Hausner, 2003), mt genomes in these organisms show signals of recombination, a characteristic that is more similar to plant mtDNAs (Clark-Walker, 1992; Bernt, Braband, *et al*., 2013; Gualberto *et al*., 2014; Lavrov and Pett, 2016; Ladoukakis and Zouros, 2017). Fungi have large intergenic regions and intron characteristics that are more similar to plant mitochondrial genomes (Zhang, 1995; Gualberto *et al*., 2014). Fungal mt genomes range in size from 1.14 kb (*Spizellomyces punctatus*) to 236 kb (*Rhiczotonia solani*) with an average of 62.68 kb (Organelle Genome Resource Database of NCBI). They can be circular or linear, are usually AT enriched, and their size variation is mostly due to presence or absence of accessory genes, mobile introns and size variation in intergenic regions. Their core gene content is largely conserved, but their relative gene order is highly variable, both between and within the major fungal phyla (Wolf and Giudice, 1988; Paquin *et al*., 1997; Aguileta *et al*., 2014). Fungal mitochondria have introns and intronic open reading frames (ORFs) classified as group I and group II introns, which differ in their sequence, structure and splicing mechanisms (Kawano, Takano and Kuroiwa, 1995; Burger, Gray and Lang, 2003; Hausner, 2003; Franz Lang, Laforest and Burger, 2007; Alverson *et al*., 2010; Bernt, Braband, *et al*., 2013). Typically, group-II introns contain ORFs that code for reverse-transcriptase-like proteins. In contrast, group-I introns can encode proteins with maturase and/or endonuclease activity (Hausner, 2003). Because of the limited comparative analysis of complete fungal mt genome sequences, it has been difficult to estimate the timeframes and molecular evolution associated with mt genes or genomes (Torriani *et al*., 2014).

The high copy number of mitochondrial genomes per cell facilitates the recovery of complete sequences by high-throughput sequencing technologies. Also, fungal inheritance of mitochondria is mostly uniparental and characterized by non-random segregation (Griffiths, 1995; Basse, 2010). Because of the limited availability of complete mt genome sequences from fungal pathogen species, it has been difficult to estimate the timeframes associated with mt genome evolution and intron fixation or invasion of introns into mt genes or genomes. Comparative analysis of nuclear genomes from Mycospharellaceae and particularly the Sigatoka disease complex already provided new insights into the evolutionary mechanisms associated with nuclear genome evolution (Chang *et al*., 2016). Comparisons between nuclear and mitochondrial genomes are useful to identify hybridization, introgression or incomplete lineage sorting among other evolutionary events (Depotter *et al*., 2018; Gonthier *et al*., 2018). A series of reports of emerging fungal hybrids, all involving plant pathogens was reviewed by Brasier (2000). New fungal hybrids are reported on a regular basis (Depotter *et al*., 2018; Giordano *et al*., 2018). Thus, investigating the evolutionary history of both nuclear and mitochondrial genomes is a matter of interest for fungal pathogens. Up until now no studies compared the mitochondrial genomes of Mycosphaerellaceae species using a timeframe.

Mycosphaerellaceae is a highly diverse fungal family containing endophytes, saprobes, epiphytes, fungicolous and phytopathogenic species in more than 56 genera (Wijayawardene *et al*., 2014; Videira *et al*., 2017). Family members can cause significant economic loss in a large number of important plants including ornamentals, food crops and commercially propagated trees. For instance, *Sphaerulina musiva* and *Sphaerulina populicola* are poplar pathogens, *Cladosporium fulvum* infects tomatoes, *Dothistroma septosporum* infects pines, and *Zymoseptoria tritici* infects wheat (Carlier *et al*., 2000; Thomma *et al*., 2005; Stukenbrock *et al*., 2006; De Lapeyre *et al*., 2010; Drenkhan *et al*., 2014; Dhillon *et al*., 2015). Three Mycosphaerellaceae members, *Pseudocercospora eumusae*, *Pseudocercospora fijiensis*, and *Pseudocercospora musae* (Wijayawardene *et al*., 2014) are major pathogens of bananas and plantains. They comprise the so-called Sigatoka disease complex which are responsible for the most economically destructive diseases for banana growers (Arzanlou, Groenewald, Fullerton, Abeln, Carlier, M.-F. Zapater, *et al*., 2008; Chang *et al*., 2016).

Diseases caused by these three pathogens induce plant physiological alterations including a reduction in photosynthetic capacity, crop yield, and fruit quality (Arzanlou, Groenewald, Fullerton, Abeln, Carlier, M. Zapater, *et al*., 2008). Symptoms are very similar for all species: *P. musae* causes yellow Sigatoka, producing mild symptoms in the form of necrotic yellow spots in leaves; *P. fijiensis* causes black Sigatoka which is more agressive and produces black necrotic leaf lesions, and *P. eumusae* disease is even more severe than black Sigatoka but induces very similar symptoms. Despite such differences in virulence, the three species share a common hemi-biotrophic lifestyle and disease cycle on their host (Chang *et al*., 2016). Sigatoka disease complex causal agents formed a robust clade, with *P. fijiensis* diverging earlier (39.9 – 30.6 MYA) than the sister species *P. eumusae* and *P. musae* (22.6 – 17.4 MYA) (Ohm *et al*., 2012; Arango Isaza *et al*., 2016; Chang *et al*., 2016). Evolutionary mechanims of mt genome evolution in the Sigatoka disease complex could be better understood using fossil calibrated phylogenies and relaxed clocks for comparative analysis of these pathogens.

Fungal plant pathogens can rapidly develop molecular mechanisms of resistance to commercially available fungicides such as the case of resistance to *QoI* fungicides by punctual mutations in the cytochrome b mitochondrial gene (Sierotzki *et al*., 2000; Ma and Michailides, 2005; Cañas-Gutiérrez *et al*., 2006, 2009).Therefore, increasing the number of fungicide applications necessary to control the disease (Cools and Fraaije, 2013; Kitchen *et al*., 2016). The extensive use of chemical fungicides increase the production cost and is harmful to both environment and crop workers (De Lapeyre *et al*., 2010; Churchill, 2011). Research related to the pathogen vital functions, such as energy production, must be intensified in order to develop new and more effective disease control strategies for food safety, human health and environment.

Here, we sequenced and annotated mitochondrial genomes of selected Mycosphaerellaceae species that diverged ∼66 MYA. We performed a comparative analysis of mitochondrial genomes of *Pseudocercospora eumusae, Pseudocercospora fijiensis,* and *Pseudocercospora musae* (causal agents of Sigatoka); *Sphaerulina musiva* and *Sphaerulina populicola* (causal agents of leaf spot and canker diseases in poplar); *Zasmidium cellare* (ethanol metabolizing, wine cellar mold) and *Zymoseptoria tritici* (wheat plant pathogen causing Septoria leaf blotch), based on its gene organization and genetic content. Comparative analyses showed that the major sources of mitochondrial genome size variation were differences in the number and size of introns in the core mt protein-encoding genes and differences in content of free standing and intronic Homing Endonuclease Genes (HEGs), hypothetical proteins and accessory genes. Although the functions and evolutionary origins of HEGs and accessory genes are unclear, their presence illustrates the plasticity of fungal mt genomes (Torriani *et al*., 2014). No evidence of incongruences was found between previously published nuclear phylogenies and our mt topology. New fossil calibrations for the Sigatoka’s complex species and mt comparative analysis aid in our understanding of the tempo and mode of evolution of these plant fungal pathogens.

## 2. Methodology

### 2.1. Sequence sources and assemblies

Eleven mitochondrial genome sequences were used for this study; seven belonging to Mycosphaerellaceae (*Pseudocercospora fijiensis*, *Pseudocercospora eumusae, Pseudocercospora musae*, *Sphaerulina musiva, Sphaerulina populicola, Zasmidium cellare,* and *Zymoseptoria tritici*); and outgroups, one from another family in Capnodiales: *Pseudovirgaria hyperparasitica)* and three from Pleosporales, the sister group of Capnodiales: *Didymella pinodes (syn. Mycosphaerella pinodes), Parastagonospora nodorum* (*syn. Phaeosphaeria nodorum*), and *Shiraia bambusicola* (Table1). Draft mitochondrial genomes were obtained either from our own sequencing data, or sequence data available in GenBank (Benson *et al*., 1999), RefSeq (O’Leary *et al*., 2016) or the MycoCosm web portal (Grigoriev *et al*., 2014). Authors, seq ID and databases consulted for obtaining each genome are listed in Table 1.

**Table 1.** Sources of genomic data used in this study.

| Species | Sequence source | Citation | Isolate | Database identifiers (genome assembly IDs) | Collection dates and locations |
| --- | --- | --- | --- | --- | --- |
| *Pseudocercospora fijiensis* | Newly sequenced | NA | 081012 | MK754071 | Colombia, Carepa, not available |
| *Pseudocercospora fijiensis*<sup>1</sup> | MycoCosm web portal | Arango Isaza et al. 2016 | CIRAD86 | Mycfi2 | Cameroon, 1988 |
| *Pseudocercospora musae*<sup>2</sup> | GenBank | Chang et al., 2016 | CBS116634 | LFZO01000000 | Cuba, not available |
| *Pseudocercospora eumusae*<sup>2</sup> | GenBank |  | CBS114824 | LFZN01000000 | France, not available |
| *Sphaerulina musiva*<sup>1</sup> | MycoCosm web portal | Dhillon et al. 2015; Ohm et al. 2012 | SO2202/ CBS133246 | Sepmu1 | Canada, Saint-Ours, QC, 2002 |
| *Sphaerulina populicola*<sup>1</sup> | MycoCosm web portal |  | p0202b/ CBS133247 | Seppo1 | Canada, Saint-Ours, QC, 2002 |
| *Zasmidium cellare*<sup>3</sup> | GenBank | Goodwin et al. 2016 | ATCC 36951/ CBS 146.36 | KX011536.1 | Not available |
| *Zymoseptoria tritici*<sup>3</sup> | RefSeq | Torriani et al. 2008 | IPO323 | NC_010222.1 | The Netherlands, Brabant, 1981 |
| *Pseudovirgaria hyperparasitica*<sup>1</sup> | MycoCosm web portal | Crous, 2014 | CBS 121739 | Psehy1 | Not available |
| *Parastagonospora nodorum*<sup>3</sup> | RefSeq | Hane et al. 2007 | SN15 | NC_009746.1 | Not available |
| *Dydimella pinodes*<sup>3</sup> | GenBank | Okorski et al. 2016 | 165/T | KT946597.1 | Poland, Tomaszkowo, not available |
| *Sharaia bambusicola*<sup>3</sup> | RefSeq | Shen et al. 2015 | zzz816 | NC_026869.1 | China, Guilin, not available |

Data type: 1. Mitochondrial contig filtered from whole genome assembly contigs, 2. Whole genome assembly contigs, and 3. Fully annotated mitochondrial genome sequence

We sequenced a total DNA sample of *Pseudocercospora fijiensis* isolate 081012 from Carepa, Antioquia, Colombia at North Carolina University using 100bp paired-end reads and Hiseq 2500 system® (Illumina, Albany, New York). Read quality was assessed by FastQC (Andrews S., 2010) and low quality reads and/or bases were trimmed using Trimmomatic (Bolger, Lohse and Usadel, 2014). High quality reads were assembled using Spades (Bankevich *et al*., 2012) at different k-mer sizes (k = 61, 71, 81, and 91) and an assembly with the highest N50 value and assembly size was selected. The chosen assembly was scaffolded by SSPACE (Boetzer *et al*., 2011), remaining gaps in the scaffolds were closed by GapFiller (Nadalin, Vezzi and Policriti, 2012) and a final genome assembly was evaluated by REAPR (Hunt *et al*., 2013). Recovered contigs were then mapped to *P. fijiensis* mitochondrial genome available in MycoCosm (Grigoriev *et al*., 2014) using Geneious 9.1.5 (Kearse *et al*., 2012). Draft mitochondrial genome sequence was deposited in GenBank (accession number: MK754071).

Mitochondrial contigs for *Pseudocercospora musae*, *P. eumusae* and *Sphaerulina populicola* were filtered from whole genome *de novo* assembled contigs (Dhillon *et al*., 2015; Chang *et al*., 2016). A BLASTn (Camacho *et al*., 2009) was ran using a BLAST database generated with the list of assembled contigs, and using as query a multi-FASTA with Electron Transport Chain genes (ETC) from published mitochondrial genomes. A reference based assembly was also performed using previously assembled contigs, and whole genomes as queries (Table 1) and ETC genes of previously published mitogenomes as reference. Contigs recovered with the above-mentioned approaches where reassembled. Reference based assembly, and *de novo* assembly were performed using the Geneious software (Kearse *et al*., 2012). An iterative mapping approach of clean reads of *Sphaerulina musiva* mitochondrial genome (SRA: SRR3927043) was performed against a reference *cox1* to generate an assembly of the mitochondrial genome. Not annotated drafts of mitochondrial genomes of *Pseudocercospora fijiensis, P. eumusae, P. musae, S. Populicola* and *Pseudovirgaria hyperparasitica* were directly downloaded from databases (Table 1).

### 2.2. Annotation

Draft mitochondrial genomes of *Pseudocercospora fijiensis, P. eumusae, P. musae, Sphaerulina populicola, S. musiva* and *Pseudovirgaria hyperparasitica* were annotated using a combination of programs as follows: Predicted ORFs were determined with a translation code for “mold mitochondrial genomes” using Geneious 9.1.5 (Kearse *et al*., 2012). Genes were identified with BLASTP (Johnson *et al*., 2008) against the non-redundant protein database at NCBI and with the MITOS web server (Bernt, Donath, *et al*., 2013). Protein domains and sequence patterns were searched using PFAM (Finn *et al*., 2014) and PANTHER (Mi *et al*., 2017). Annotations obtained for the aforementioned approaches were contrasted and inconsistencies regarding length and position of genes were solved through multiple alignment of related gene sequences using MUSCLE (Edgar, 2004) and CLUSTALW (Larkin *et al*., 2007). We considered as hypothetical proteins those ORFs larger than 300bp that did not show results with the above-mentioned annotation strategies.

### 2.3. PCR amplifications of *cox1* gene copies in *P. fijiensis*

A PCR assay to confirm the presence of three different copies for *cox1* in *P. fijiensis* mitochondrial genome was performed. Primers were designed to amplify regions between *cox1_1* and *cox1_2*, between *cox1_2* and *cox1_3* and between *cox1_1* and *cox1_3* (Figure 1). PCR amplifications were carried out in a total volume of 10μl, containing 20ng genomic DNA, 0.15μM of each primer, 1x PCR buffer (without MgCl_2_), 0.75mM MgCl_2_, 4μM of each dNTP and 0.65 U recombinant Taq DNA polymerase (ThermoFisher, MA, USA). Cycling parameters were: 3 min at 94°C, followed by 35 cycles of 30s at 94°C, 30s at a 50 to 60 °C temperature gradient to determine annealing temperature, 1 min at 72 °C, and a final elongation step of 5 min at 72 °C. PCR products were separated by electrophoresis in a 1 % (w/v) agarose gel and visualized with GelRed® under UV light.

**Figure 1.**
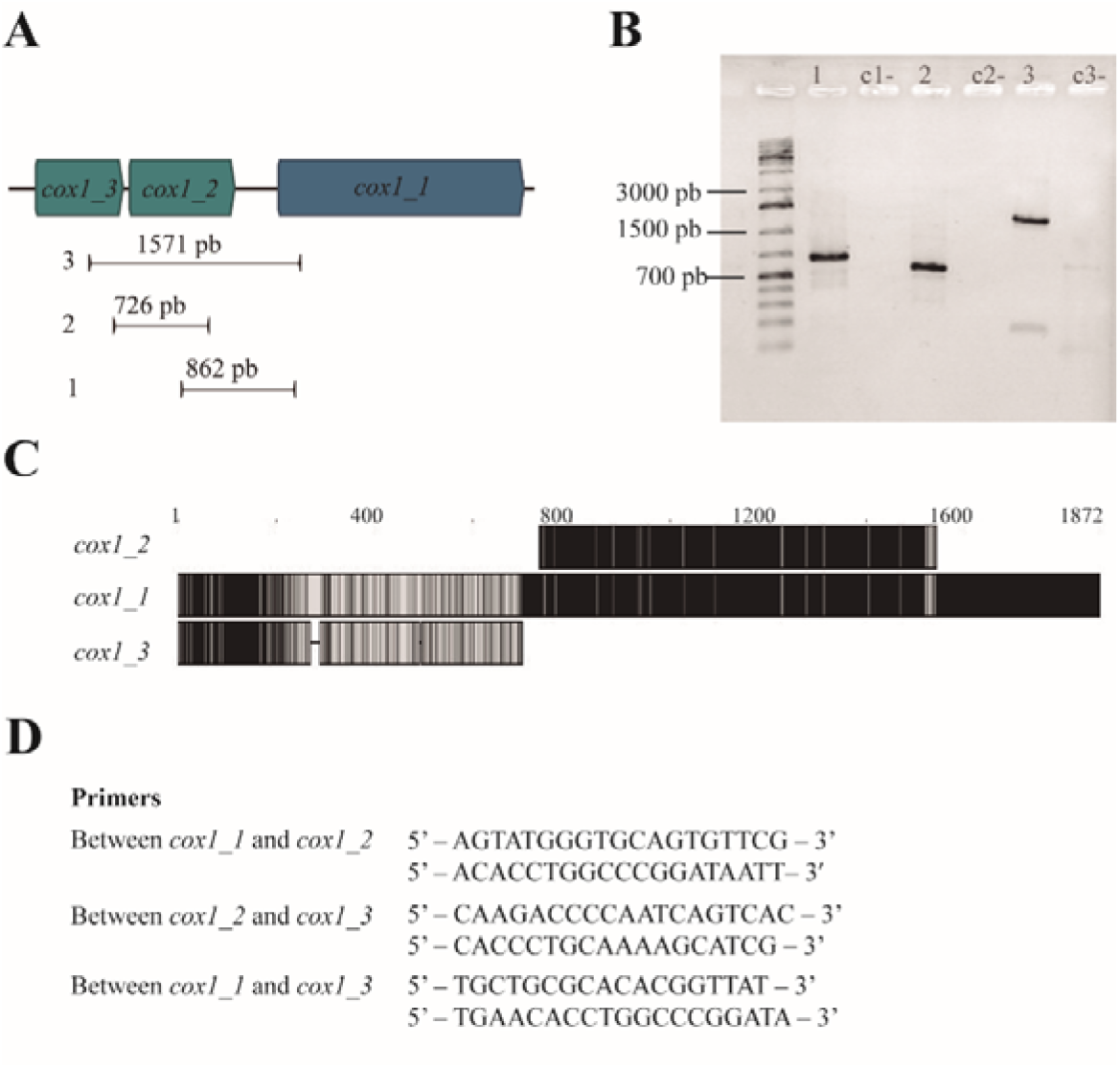
Mitochondrial gene duplication in *Pseudocercospora fijensis*. A. The three *cox1* copies are colinearly localized in the genome. B. Amplification of intergenic sequences among the copies confirmed their existence C. The copies have different size and are not identical en sequence. Identical sites apears in black in the alignment.

### 2.4. Transcriptome de novo assembly

RNA seq raw reads of *S. musiva* (SRR1652271) and *P. fijiensis* (SRR3593877, SRR3593879) were downloaded from EMBL EBI database. Reads were quality filtered and trimmed using BBDuck from the BBtools package (https://jgi.doe.gov/data-and-tools/bbtools/) before carrying out transcriptome *de novo* assemblies using the short-read assembly program “Trinity” (Grabherr *et al*., 2011). The *P. eumusae* (GDIK00000000.1) and *P. musae* assembled transcriptomes (GDIN00000000.1) were downloaded from GeneBank. We mapped transcripts to assembled genomes and genes of interest using Geneious mapper for RNA seq.

### 2.5. RT –PCR assays for mitochondrial gene pairs of *P. fijiensis*

Total RNA was extracted from *P. fijiensis* mycelium after fifteen days of culture using TRIzol® reagent (Life Technologies) according to the manufacturer’s instructions. RNA concentration was measured at 260 nm, using a NanoDrop ND-1000 UV-Vis Spectrophotometer (NanoDrop Technologies, Thermo Fisher). DNAse I (Thermo Scientific) was used to treat RNA and cDNA molecules were synthetized using Maxima First Strand cDNA Synthesis Kit (Thermo Scientific) according to manufacturer’s instructions and used as template for amplification. Primers were designed such that the amplified product encompass the end of one gene and the beginning the other for gene pairs *nad3-nad2* and *atp8-atp6* and amplifying partially both genes and their intergenic sequences. PCR amplification was performed and PCR products were analyzed by 1% agarose gel electrophoresis and gel purified using the GFX PCR DNA and gel band purification kit, according to the manufacturer’s instructions (GE Healthcare). All purified PCR products were sequenced by Sanger Technology at Macrogen Inc (http://www.macrogen.com).

### 2.6. Phylogenetic analysis and divergence times estimates

To analyze the evolution of plant fungal pathogens through time and across sampled species, we first reconstructed a phylogenetic tree, then calculated reliable diversification times and last mapped mt molecular features such as size, gene order and others, in an evolutionary context. Since most mt genomes are uniparentally inherited we used our mt phylogeny to compare topologies with already published nuclear ones for the Sigatoka disease complex species (Arango Isaza et al. 2016; Chang et al. 2016). Up until now only nuclear markers and strict clock calibration have been used to calculate diversification times for the Sigatoka disease complex species. We aim to compare these analyses with fossil calibrated Bayesian phylogenies and mitochondrial markers.

To reconstruct concatenated gene trees, ten out of fourteen core mitochondrial genes (*cox1, cox3, atp6, cob, nad1, nad2, nad4, nad4L, nad5, nad6*) belonging to eleven Mycospharellaceae species were aligned using CLUSTALW (Larkin *et al*., 2007). A Generalized time reversible (GTR) model was used with an estimated gamma parameter of rate heterogeneity to build maximum likelihood (ML) trees using the Randomized Accelerated Maximum Likelihood (RAxML) (Stamatakis, 2014) and PhyML (Guindon *et al*., 2010) programs. One hundred bootstrapped trees were generated and used to assign bootstrap support values to the consensus trees. The species *Didymella pinodes* (Okorski *et al*., 2016), *Pseudovirgaria hyperparasítica* (Braun *et al*., 2011)*, Phaeosphaeria nodorum* (Hane *et al*., 2007) and *Shiraia bambusicola* (Shen *et al*., 2015) were used as outgroup.

A bayesian phylogeny and divergence time analysis were carried out using BEAST2 version 2.5.1 (Bouckaert *et al*., 2014). Separate partitions for each gene were created with BEAUti2 (BEAST 2 package). More suitable substitution models for each gene were found using the software package jModelTest2 (Guindon, Gascuel and Rannala, 2003; Darriba *et al*., 2012) according to the Bayesian Information criterion (BIC) (Schwarz, 1978). To accommodate for rate heterogeneity across the branches of the tree we used an uncorrelated relaxed clock model (Drummond *et al*., 2006) with a lognormal distribution of rates for each gene estimated during the analyses. We also used a strict clock to further compare results. The fossil Metacapnodiaceae (Schmidt *et al*., 2014) was used assuming to be a common ancestor of the order Capnodiales with minimum age of 100 MYA (gamma distribution, offset 100, mean 180, maximum soft bound 400). Capnodiales nodes were constrained to monophyly based on the results obtained from ML analysis. A birth/death tree prior was used to model the speciation of nodes in the topology, with gamma priors on probability of splits and extinctions. We used vague priors on the substitution rates for each gene (gamma distribution with mean 0.2 in units of substitutions per site per time unit). To ensure congruence we ran the analyses five times for 50 million generations each, sampling parameters every 5000 generations, assessing convergence and sufficient chain mixing (Effective sample sizes > 200) using Tracer 1.5 (Rambaut and Drummond, 2009). After removal of 20 % of each run as burn in the remaining trees were combined using LogCombiner (part of the BEAST package), summarized as maximum clade credibility (MCC) trees in TreeAnnotator (part of the BEAST package), and visualized using FigTree. (Rambaut, 2006)

## 3. Results

### 3.1. Inferred mt phylogeny and diversification times

A comparative framework was used to understand mitochondrial evolution in fungal plant pathogens. Since most mt genomes are uniparentally inherited we built a mt phylogeny in order to compare topologies with already published nuclear ones for the Sigatoka disease complex species. Our mitochondrial topology supports previously published nuclear phylogenies (Arango Isaza et al. 2016; Chang et al. 2016) and all nodes were well supported (Figure 6). Ideally, phylogenies should be built with single-copy genes to avoid paralogy (Guindon, Gascuel and Rannala, 2003). However, most core mitochondrial genes for species included in this phylogeny have more than one copy. ML phylogenies were built for each gene including all copies per specie. All gene copies were monophyletic in all cases (Appendix A3-14), indicating that duplicated genes are still poorly differentiated and homology among copies from same species can still be detected. One complete copy with start and stop codons was selected to build a species tree. A phylogeny using only a single copy *rns* gene was also used. The same topology was always recovered (Appendix A15). Finally, we used ten of fourteen core genes for divergence time estimation because when using only the *rns* gene, the 95% highest posterior density (HPD) intervals were significantly broader.

A robust phylogeny with posterior probabilities greater than 0.97 was recovered, containing two main linages (Pleosporales + Capnodiales). Divergence time estimates using fossil calibration are shown in Figure 6, with horizontal bars representing the 95% highest posterior density (HPD) intervals for each node. According to our data, Mycosphaerellaceae diverged from the rest of Capnodiales at the end of the Mesozoic or the early Paleogene, about 66.66 MYA (55.47 −78.27 MYA, 95% HPD). The earliest split within Mycosphaerellaceae gave rise to *Zasmidium cellare*. The species *Zymoseptoria tritici* diverged from (*Sphaerulina* + *Pseudocercospora*) also at the end of the Mesozoic or the early Paleogene 59.88 MYA (49.7-70.91 95% HPD). The sister genera *Sphaerulina* + *Pseudocercospora* diverged in the Eocene, 48.1 MYA (39.34-57.86, 95% HPD); while the sister clade to the species in the Sigatoka complex includes the species *Sphaerulina musiva* and *S. populicola* sharing their last common ancestor during the Miocene, 13.39 MYA (9.39-17.69 MYA, 95% HPD). The origin of *Pseudocercospora* members of the Sigatoka disease complex in banana was dated around 13.31 MYA (9. 49-17.28, 95% HPD) during the Miocene while the sister *species P. eumusae* and *P. musae* split from their shared ancestor in the late Miocene 8.22 MYA (5.6 −11.07, 95% HPD) (Figure 6).

### 3.2. Mitochondrial genome content and genome organization

We compared mitochondrial genomes of *Pseudocercospora fijiensis, P. eumusae, P. musae, Sphaerulina populicola,* and *S. musiva*, with previously published mitochondrial genomes of *Zymoseptoria tritici* and *Zasmidium cellare.* They showed a homogeneous GC content between 27% and 32%, high variability in genome size, ranging from 23743 bp in “cellar mold” *Zasmidium cellare* to 136906 bp in poplar pathogen *Sphaerulina populicola* (Appendix 1). Annotated genes of seven mitochondrial genomes analyzed included a set of conserved mitochondrial protein coding genes, namely the subunits of the electron transport chain (ETC) complex I (*nad1*, *nad2*, *nad3*, *nad4*, *nad4L*, *nad5* and *nad6*), complex III (*cox1*, *cox2* and *cox3*), complex IV (*cob*) and ATP synthase subunits (*atp6*, *atp8* and *atp9*). A small ribosomal subunit RNA (rns), a large ribosomal subunit rRNA (rnl), and a set of transfer RNAs (tRNAs). Five of seven genomes contained hypothetical proteins, accessory proteins, additional copies of genes and genetic mobile elements (Table 3). ETC gene order among Mycosphaerellaceae mitochondrial genomes vary greatly, except for sister species *Sphaerulina populicola* and *S. musiva* which showed the same gene order (Figure 3). Despite this variability, gene pair order was conserved for *nad4L*-*nad5*, *nad3-nad2* and *atp8-atp6* among all mitochondrial genomes (Figure 3). We mapped RNA seq assembled transcripts to *P. fijiensis, S. musiva, P. musae* and, *P. eumusae* mitochondrial genomes and found that these neighbor genes were contained in the same transcript suggesting that they were transcribed as a single mRNA.

**Figure 2.**
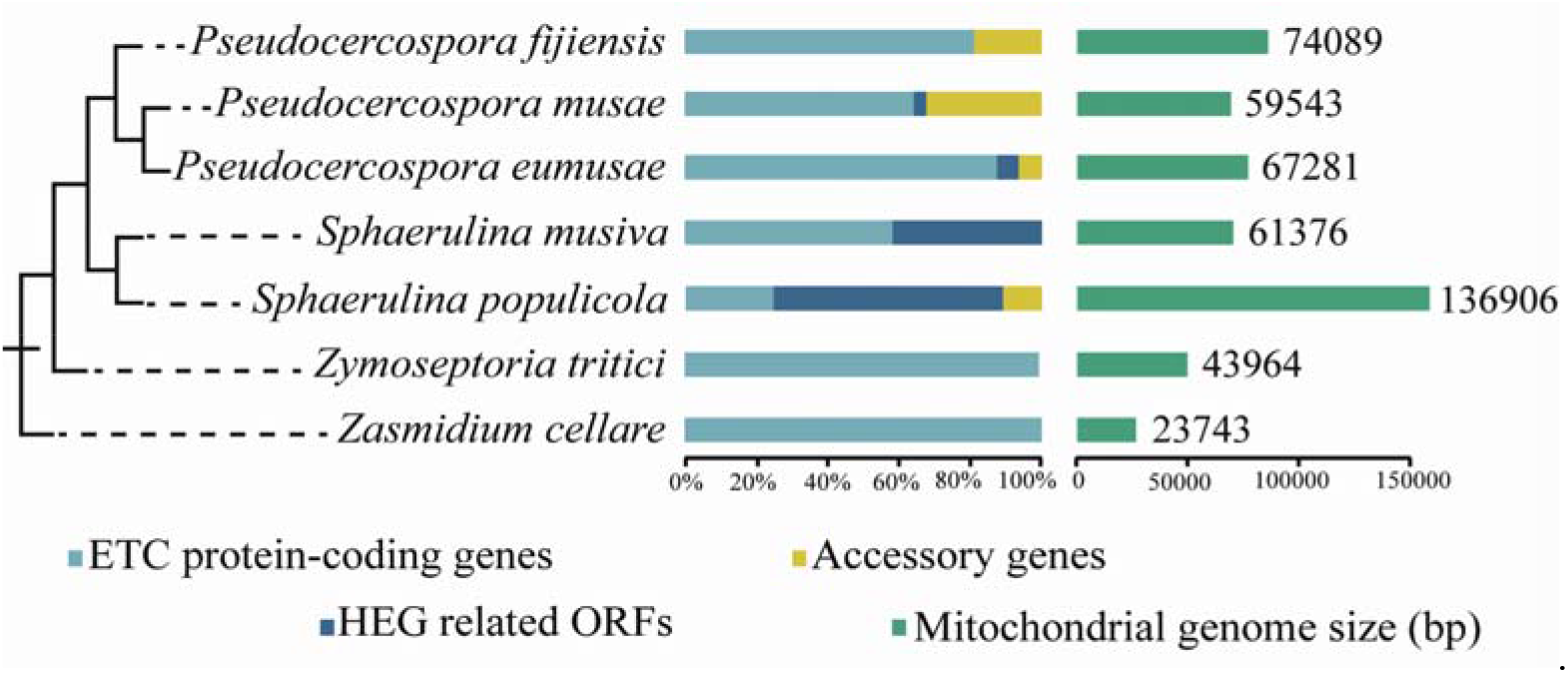
Estimated mitochondrial genome size among Mycosphaerellaceae and contribution of ETC genes, hypothetical proteins, HEG related ORFs and accessory genes to genetic content of mitochondrial genomes. Mitochondrial genomes of Mycosphaerellaceae members are diferent in terms of genome size and content of accessory genes and HEG. Some genomes contain only the ETC transport genes while others exhibit HEG invasion

**Figure 3.**
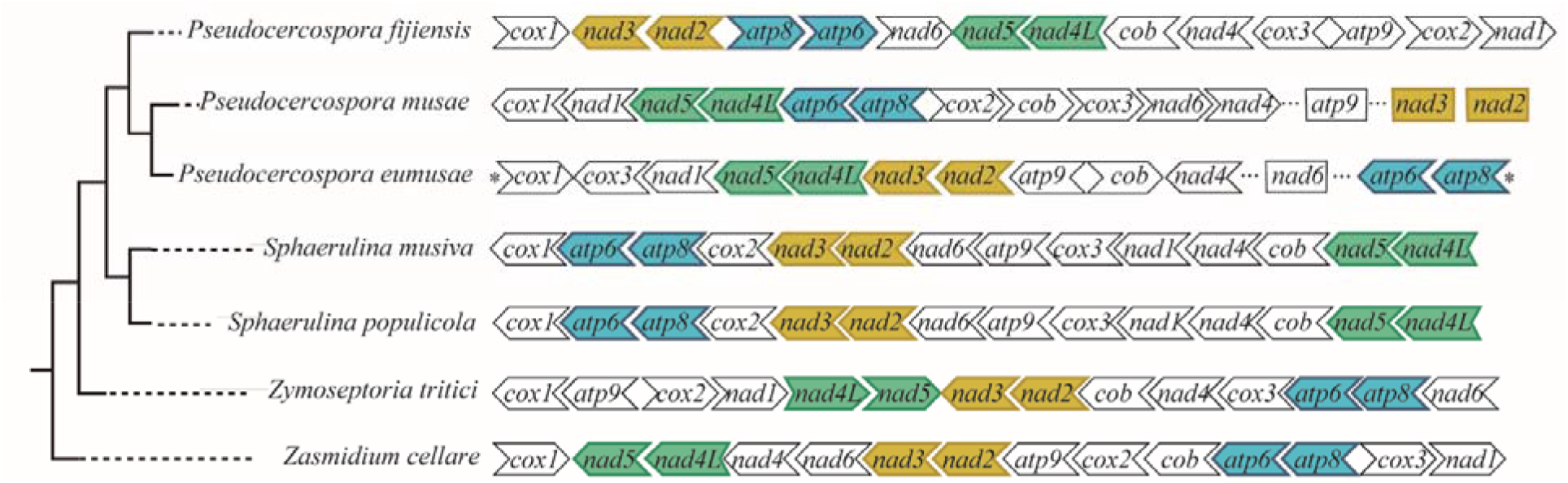
Core genes order and orientation mapped to Mycosphaerellaceae recovered mitochondrial phylogeny. Gene order is not conserved across members except for *Sphaerulina* species. Colored gene pairs were always recovered as neighbors, asterisks represent continuity of the sequences.

All annotated genomes were found to have duplications of some ETC genes. They were considered as gene duplications because they presented the same PFAM and PANTHER domains and high local sequence similarity (>90%). Duplicated gene alignments showed that gene copies were not identical in length (Figure 1C), and only one copy had a complete coding sequence. The mitochondrial genome of *P. fijiensis* has the greater number of duplicated genes, with five ETC genes having two to three copies (Tables 2 - 3). Duplicated genes were found either close to each other, or disperse in the genomes.

### 3.3. Experimental confirmation of gene copies, bicistronic transcription and intron content in *P. fijiensis* mt genome

We used *P. fijiensis* to prove experimentally the three most characteristic features of Mycosphaerellacea genomes found through bioinformatics analysis: Presence of ETC copies, bicistronic transcription of some genes and intron presence in ETC genes. First, PCR amplifications were performed to confirm the presence of three *cox 1* copies in *P. fijiensis* mitochondrial genome. Primers were designed to amplify regions among the three different copies of *cox1* (Figure 1D). We chose to amplify this region because it was the only case of tandem duplication among all genomes. Other gene copies in *P. fijiensis* as well as in other species were found sparse. The three regions amplified with bands of the expected size, confirming that the different gene copies exist (Figure 1A and B). Second, in *P. fijiensis* mitochondrial genome there are two genes with introns (*cob* and *nad5*), so we confirmed the presence of *cob* intron experimentally but amplification of *nad5* intron was unsuccessful because the PCR product exceeds 3Kb and there were few optimal primer design sites surrounding the targeted region (Appendix 2). Finally, RT-PCR amplification and subsequent Sanger sequencing confirmed polycistronic expression for *nad3-nad2*, *atp8-atp6* gene pairs in *P. fijiensis* (Figures 3-4).

**Figure 4.**
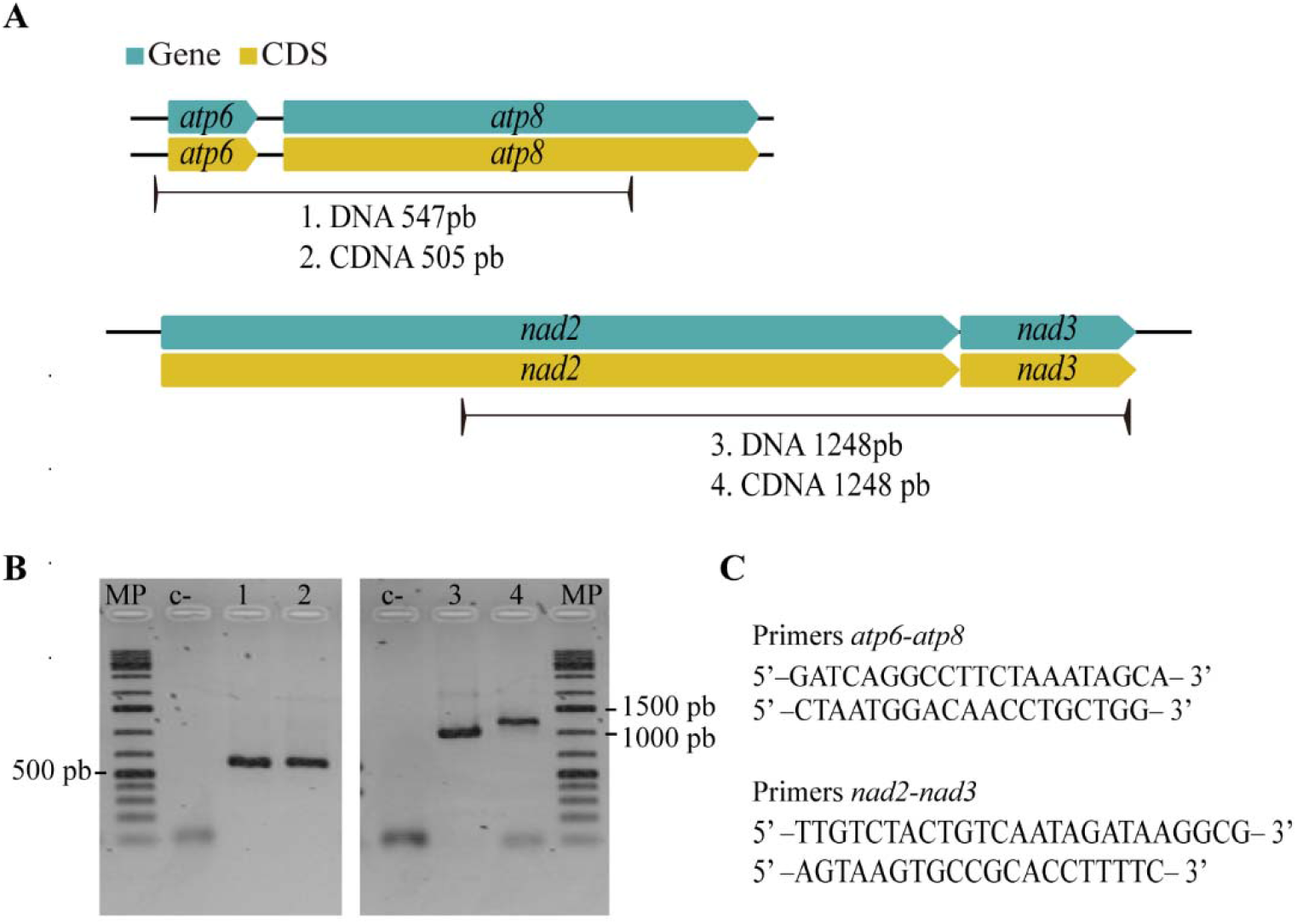
RT-PCR assay to confirm bicistronic expresion of genes. Designed primers covering gene pairs *nad2-nad3*, *atp6-atp8* and its intergenic sequences amplifiyed bands of the expected.

### 3.4. Homing endonucleases invasion and putative horizontal gene transfer

Differences in genome sizes among members of the Mycosphaerellaceae were attributed to the presence of different amounts of accessory genes, gene copies and intron mobile sequences (Figure 2, Tables 2-3). Accessory genes, gene copies and intron mobile sequences are specie specific. A commonly observed feature in Mycosphaerellaceae mitochondrial genomes was an invasive presence of sequences containing LAGLIDADG or GIY-YIG domains related to homing endonuclease genes (HEG) (Chevalier and Stoddard, 2001). These sequences caused fragmentation in ETC genes. In almost all instances, pieces of an ETC genes were collinearly distributed and each fragment was found to be followed by an insertion of an HEG related sequence. An extreme case was found in *cox1* in *Sphaerulina populicola* (CDS: 1542bp), which has twelve fragments distributed along 22070 bp, each of them containing a piece of *cox1* followed by an HEG related domain (Figure 5).

**Figure 5.**
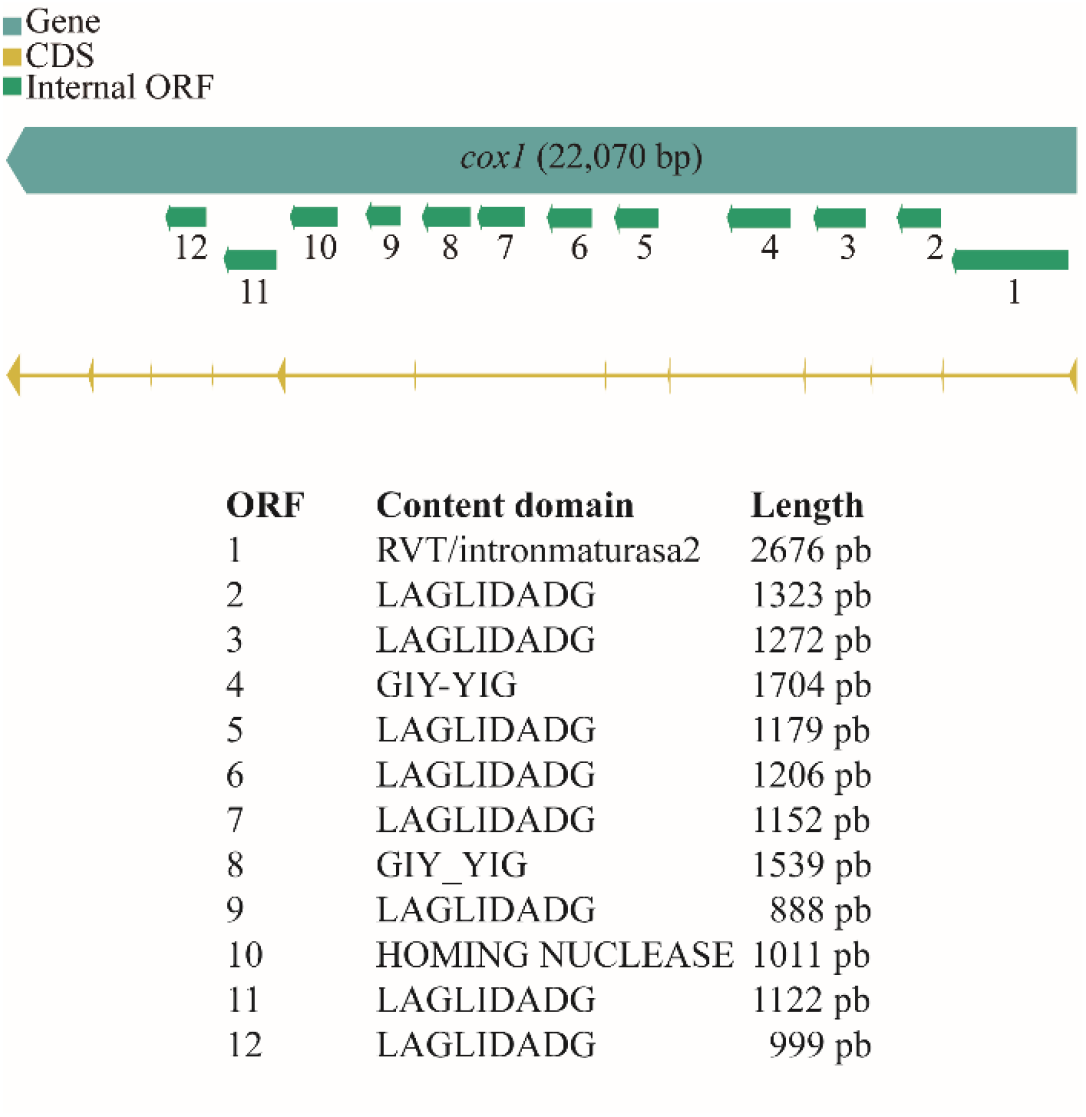
Homing endonuclease invasion in the mitochondrial genome of *Sphaerulina populicola*. HEG invasion in *cox1* fragmented the gene into twelve exons. Numbers 1-14 represent different internal open reading frames (ORF). Domains content in the ORFs are listed in the table along with their position within *cox1*.

**Figure 6.**
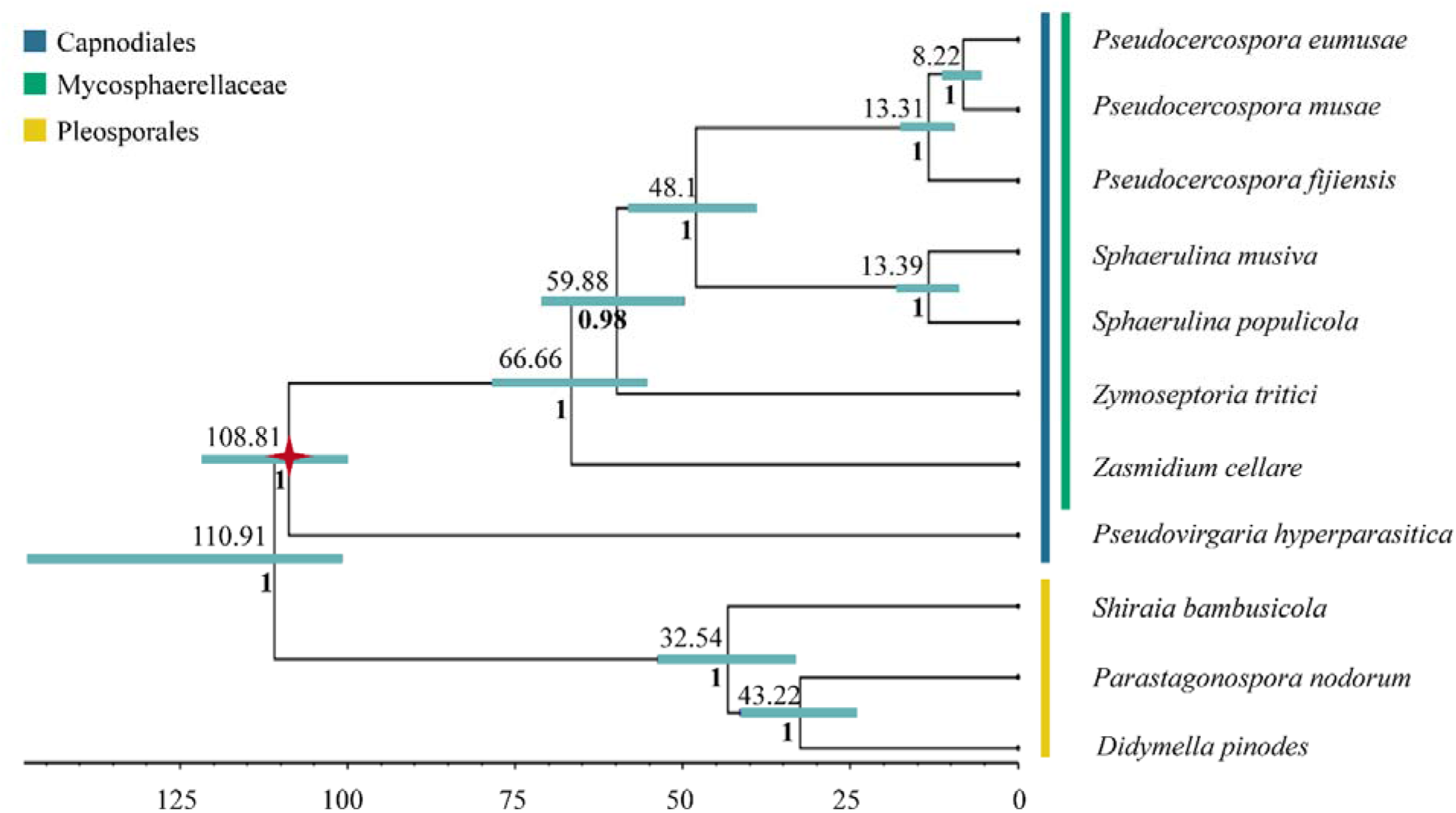
Mycosphaerellaceae bayesian phylogeny and divergence time estimation. Mycosphaerellaceae bayesian phylogeny with posterior probabilities in each branch and divergence time estimates calculated using fossil calibrations in BEAST2. Red star represents the fossil calibration placement.

## 4. Discussion

Our principal objective was to investigate the evolutionary history of mitochondrial genome evolution in fungal plant pathogens such Mycosphaerellaceae. We found unequivocal evidence of at least five independent horizontal gene transfer events related to variation in genome composition in this family. Even though we found variation in mitochondrial genome composition there was no evidence of hybridization when comparing nuclear and mitochondrial dataset sets for fungal plant pathogens analyzed here.

Genome variation in size and organization is not correlated with phylogenetic relationships, nor does it carry phylogenetic signal based on our current sampling. Ascomycete’s mitochondrial genomes like most mitochondrial genomes along the tree of life generally consist of a highly conserved core of protein encoding genes: two ribosomal subunits (*rnl* and *rns*), a distinct set of tRNAs and fourteen genes of the respiratory chain complexes (*cox1*, *cox2*, *cox3*, *cob*, *nad1* to *nad6*, *atp6*, *atp8* and *atp9*) (Clark-Walker, 1992). These were also the genes found in mitochondrial genomes of Mycosphaerellaceae. Besides this common set of genes, a variable number of free standing open reading frames of unknown function and a variable number of introns related with homing endonucleases (HEF), often including GIY-YIG or LAGLIDADG protein domains were found. These ORFs ranged from one in *Pseudocercospora musae* and *P. eumusae* to 36 in *Sphaerulina populicola* (Table 2). Despite not being ubiquitous in Ascomycete mitochondrial genomes, these ORFs are common (Clark-Walker, 1992; Adams and Palmer, 2003; Burger, Gray and Lang, 2003; Hausner, 2003; Lodish, 2003; Bernt, Braband, *et al*., 2013; Jelen *et al*., 2016). HEG invasion has been previously described in fungi, where fragmented genes are the rule (Dassa *et al*., 2009). However, it is not common to find highly fragmented genes as the *cox1* of *Sphaerulina populicola* (Figure 5). A similar case was reported in the mitochondrial genome of *Sclerotinia borealis*, where researchers found thirteen introns of *cox1* and truncated copies of ETC genes (Mardanov *et al*., 2014).

**Tabla 2.** Mycosphaerellaceae mitochondrial genome molecular features.

|  | *P. musae* | *P. eumusae* | *P.*<br>*fijiensis* | *S. musiva* | *S.*<br>*populicola* | *Z. cellare* | *Z. tritici* |
| --- | --- | --- | --- | --- | --- | --- | --- |
| **Estimated mitochondrial genome size (bp)** | 59543 | 67281 | 74089 | 61376 | 136906 | 23743 | 43964 |
| **GC content (%)** | 27.80 | 27 | 26.00 | 28.80 | 31.70 | 27.80 | 31.90 |
| **Coding percentage (%)** | 69.91 | 94% | 65.55 | 51.50 | 53.99 | 84.90 | 64.56 |
| **ETC protein coding genes** | 14 | 14 | 14 | 14 | 14 | 14 | 14 |
| **Additional gene copies** | 7 | 1 | 7 | 2 | 1 | 0 | 0 |
| **Hypothetical proteins** | 8 | 3 | 12 | 5 | 8 | 1 | 0 |
| **HEG related ORFs** | 1 | 1 | 3 | 10 | 36 | 0 | 0 |
| **Accessory Genes** | 7 | 2 | 0 | 0 | 6 | 0 | 0 |
| **tRNA** | 25 | 25 | 40 | 28 | 30 | 25 | 27 |
| **rns size bp** | 1587 | 1947 | 1925 | 1625 | 1705 | 1529 | 1476 |
| **rnl size bp** | 3493 | 3505 | 9652 | 5165 | 8280 | 3079 | 1673 |

<sup>1</sup>I.(nucleotides in protein-coding genes, rRNAs and tRNAs) /Total size.

Differences in mitochondrial genomes sizes among members of Mycosphaerellaceae are attributed to heterogeneous content of accessory genes, gene copies and intron mobile sequences in some of the genomes (Figure 2, Tables 2-3). These elements are species specific, indicating that they have been acquired through horizontal transfer through at least five independent events in their evolutionary history (Figure 5). In *Pseudocercospora* and *Sphaerulina* species we found duplicated genes, ORFs related to HEGs and hypothetical proteins, all contributing to the variation in size. Mitochondrial genomes from *P. musae*, *P. eumusae*, and *S. Populicola* had also accessory genes including RNA and DNA polymerases, reverse transcriptases and transposases. Even for sister species such as *Sphaerulina musiva* and *S. populicola* who share core gene content (ETC genes) and gene order, genomes sizes differ greatly due to a higher content of homing endonucleases HEG related ORFs and the presence of accessory genes in *S. populicola* (Appendix 1). Invasion of mobile elements and accessory genes as main source of size variability among the genomes of phylogenetically related organisms was observed before in a study where nine mitochondrial genomes of *Aspergillus* and *Penicilium* were compared (Joardar *et al*., 2012). In the phytopathogenic fungus *Sclerotinia borealis*, they also found expansion of the genome due to plasmid like sequences integrated in the mitochondrial DNA and HEG related ORFs (Mardanov *et al*., 2014).

Mitochondrial genomes of *S. musiva, S. populicola, P. fijiensis, P. eumusae* and *P. musae* have partial extra copies of ETC protein coding genes (Table 3) and *tRNAs*, while the fully annotated published Mycosphaerellaceae genomes of *Zasmidium cellare* and *Zymoseptoria tritici* do not possess any duplication (Torriani *et al*., 2008; Goodwin *et al*., 2016). Mitochondrial gene duplication is seldom described in fungi. Maradanov et al. (2014) found duplications of some mitochondrial regions, resulting in the appearance of truncated extra copies of *atp9* and *atp6* in phytopathogenic fungus *Sclerotinia borealis* (Mardanov *et al*., 2014). Two incomplete copies of *atp6* were found on different strands of the mitogenomic DNA of *Shiraia bambusicola*, a Pleosporales member used as outgroup in this study (Shen *et al*., 2015). Three hypotheses can be drawn regarding mitochondrial gene duplications: First, copies could be undergoing subfunctionalization (Conant and Wolfe, 2008) where recombination events of fission and fusion are the drivers of such a process (Burger, Gray and Lang, 2003), however we did not see two functional copies of any ETC genes in any genome.. Second, they can be truncated copies accounting for pseudogenes (Conant and Wolfe, 2008). We therefore suggest that additional Mycospharellaceae fungi need to be available before stronger conclusions can be drawn about the proximal causes of gene duplication. Further gene editing and virulence assays to test these hypotheses can shed light on fungal adaptation.

**Tabla 3.**
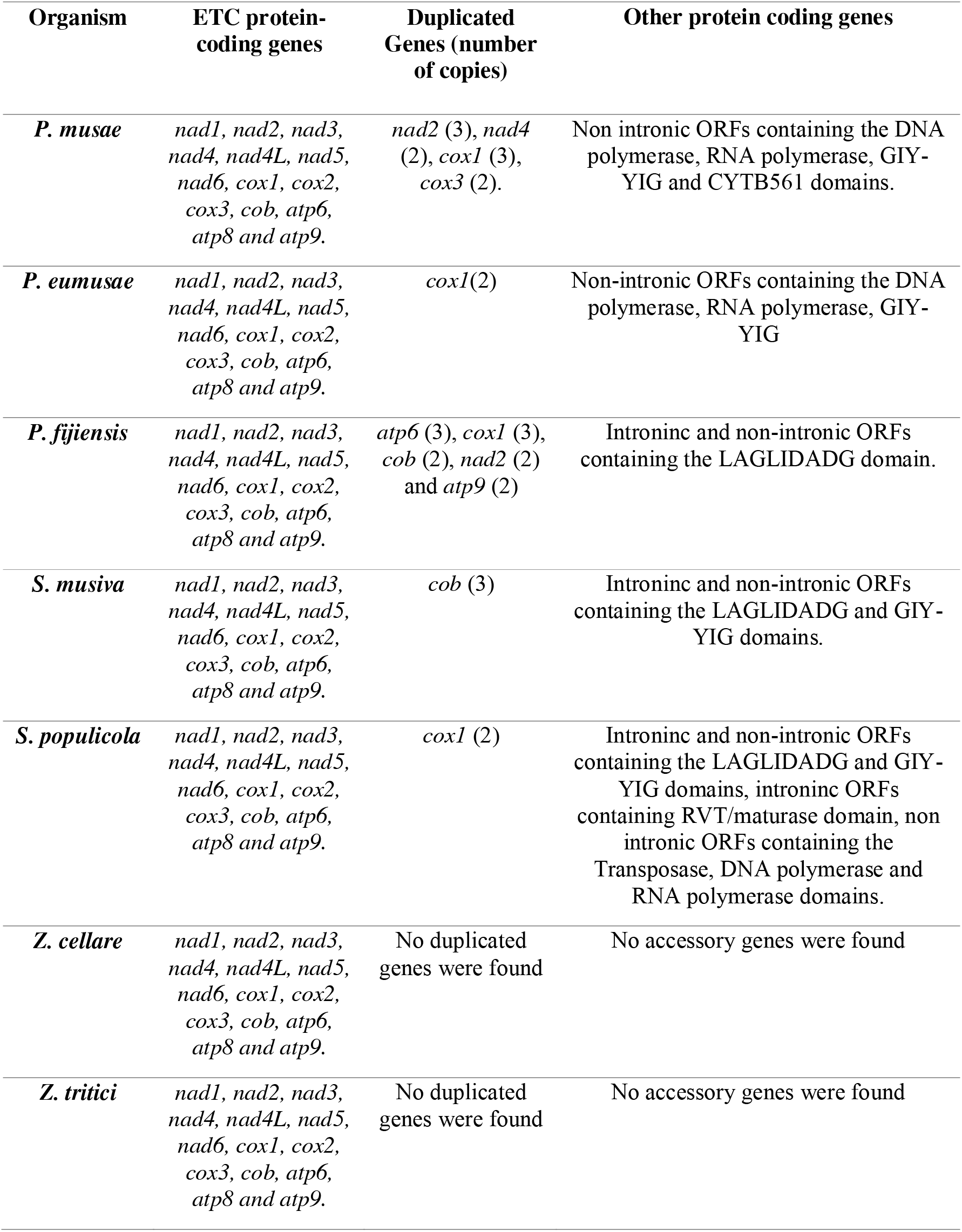
Protein-coding genes annotated in Mycosphaerellaceae mitochondrial genomes.

| Organism | ETC protein-coding genes | Duplicated Genes (number of copies) | Other protein coding genes |
| --- | --- | --- | --- |
| *P. musae* | *nad1, nad2, nad3, nad4, nad4L, nad5, nad6, cox1, cox2, cox3, cob, atp6, atp8 and atp9.* | *nad2* (3), *nad4* (2), *cox1* (3), *cox3* (2). | Non intronic ORFs containing the DNA polymerase, RNA polymerase, GIY-YIG and CYTB561 domains. |
| *P. eumusae* | *nad1, nad2, nad3, nad4, nad4L, nad5, nad6, cox1, cox2, cox3, cob, atp6, atp8 and atp9.* | *cox1*(2) | Non-intronic ORFs containing the DNA polymerase, RNA polymerase, GIY-YIG |
| *P. fijiensis* | *nad1, nad2, nad3, nad4, nad4L, nad5, nad6, cox1, cox2, cox3, cob, atp6, atp8 and atp9.* | *atp6* (3), *cox1* (3), *cob* (2), *nad2* (2) and *atp9* (2) | Introninc and non-intronic ORFs containing the LAGLIDADG domain. |
| *S. musiva* | *nad1, nad2, nad3, nad4, nad4L, nad5, nad6, cox1, cox2, cox3, cob, atp6, atp8 and atp9.* | *cob* (3) | Introninc and non-intronic ORFs containing the LAGLIDADG and GIY-YIG domains. |
| *S. populicola* | *nad1, nad2, nad3, nad4, nad4L, nad5, nad6, cox1, cox2, cox3, cob, atp6, atp8 and atp9.* | *cox1* (2) | Introninc and non-intronic ORFs containing the LAGLIDADG and GIY-YIG domains, introninc ORFs containing RVT/maturase domain, non intronic ORFs containing the Transposase, DNA polymerase and RNA polymerase domains. |
| *Z. cellare* | *nad1, nad2, nad3, nad4, nad4L, nad5, nad6, cox1, cox2, cox3, cob, atp6, atp8 and atp9.* | No duplicated genes were found | No accessory genes were found |
| *Z. tritici* | *nad1, nad2, nad3, nad4, nad4L, nad5, nad6, cox1, cox2, cox3, cob, atp6, atp8 and atp9.* | No duplicated genes were found | No accessory genes were found |

Mycosphaerellaceae gene order is not conserved among mitochondrial genomes (Figure 3). Even though divergence times within sister clades *Pseudocercospora* 13.31 MYA (9.49-17.28, 95% HPD) and *Sphaerulina* 13.39 MYA (9.39-17.69 MYA, 95% HPD) are roughly the same (Figure 6), gene order is highly conserved in the *Sphaerulina* species but not in the *Pseudocercospora* ones (Figure 2). Nonetheless, some pairs of genes are always found together *atp6-atp8, nad2-nad3, nad4L-nad5*. Here, bicistronic transcription of *atp6-atp8*, *nad2-nad3* through RT-PCR was confirmed (Figures 3-4). The presence of mitochondrial transcripts containing more than one ORF make sense due to mt endosymbiotic origin from an α-proteobacteria (Schäfer, Hansen and Lang, 2005; Barbrook *et al*., 2010; Chrzanowska-Lightowlers, 2015). We attribute differences in gene order to genomic rearrangements, which could be caused by different processes, such as fusion, fission, recombination, plasmid integration and mobility among mitochondrial genomes (Kawano, Takano and Kuroiwa, 1995). Indeed, plasmid integration appears to have a role in recombination thanks to the presence of long inverted repeats that allows homologous recombination (Salavirta *et al*., 2014). There are no broad studies comparing mitochondrial gene order among closely related species of fungi. Aguileta et al. (2014) analyzed 38 complete fungal mitochondrial genomes to investigate the evolution of mitochondrial DNA gene order among fungi. They looked for evidence of non-homologous intra chromosomal recombination and investigated the dynamics of gene rearrangements and the effect introns, intronic ORFs and repeats may have in gene order. Their results support recombination in all fungal phyla. Despite the big gene order differences found, they noticed conserved pairs of genes *nad2-nad3* and *nad4L-nad5* in most genomes (Aguileta *et al*., 2014) like our results. Maintenance of such proximity through evolution is important for mitochondria functioning and modifications of such proximity can affect negatively the viability of organisms.

Since most mt genomes are uniparentally inherited we built a mt phylogeny in order to compare topologies with already published nuclear ones for the Sigatoka disease complex species (Arango Isaza et al. 2016; Chang et al. 2016). When phylogenetic comparisons between mitochondrial and nuclear data sets in fungi differ, hybridization, introgression or incomplete lineage sorting among other events could explain incongruences (Depotter et al. 2018; Giordano et al. 2018). Our mitochondrial topology supports previously published nuclear phylogenies without indication of hybridization events. However, our calibrated phylogeny has later diversification times for the origin of Mycosphaerellaceae 66.66 MYA (55.47 −78.27 MYA, 95% HPD) compared to 186.7–143.6 MYA (Chang *et al*., 2016). This suggests that a maximum of 78 MY elapsed between the split of Mycospharellaceae species and the at least five independent intron invasion events into their mt genomes. The most parsimonious explanation for these independent intron invasions is horizontal transfer from donor species. Fungal species can directly exchange genetic material through mycelial or conidial anastomoses if they share the same ecological niche (Torriani *et al*., 2014). Another possibility is indirect transmission of genetic material through virus that can move DNA or RNA from one fungal species to the other (Rosewich and Kistler, 2000)

As far as the Sigatoka disease complex species diversification we found an early divergent *Pseudocercospora fijiensis* that splits from the sister species *P. musae* + *P. eumusae* 13.31 MYA (9.49-17.28, 95% HPD); while the sister species *P. eumusae* and *P. musae* split from their shared ancestor in the late Miocene 8.22 MYA (5.6 −11.07, 95% HPD) (Figure 6). Chang et al. (2016) using nuclear phylogenies estimated the splitting of *P. fijiensis* from their last common ancestor with *P. musae* + *P. eumusae* to be between 39.9-30.6 MYA and the splitting of *P. musae* and *P. eumusae* to be between 22.6 – 17.4 MYA. Our results naturally prompt a further question: Was the origin of the Sigatoka disease complex through host-tracking evolution? This hypothesis explain that a pathogen is likely to be younger that the host until changes related to the genetics of the pathogens or/and exogenous factors have prompted the observed alterations in their virulence spectra and the sudden flare up and over dominance of one species over the others (Stukenbrock and Mcdonald, 2008). We are currently limited in terms of a good taxonomic sampling and biogeographical analysis for *Pseudocercospora* species to answer this question.

Strong differences in diversification times compared to Chang et al. (2016) were found due to differences in calibrations points and methods. Chang et al. (2016) used r8s Likelihood methods (Sanderson, 2003) and a calibration at the origin of the Dothideomycetes crown group (394-285 MYA) using previous bayesian estimations (Gueidan *et al*., 2011). Here bayesian analysis in BEAST2 (Bouckaert *et al*., 2014) and a fossil calibration using a Metacapnodiaceae (Schmidt *et al*., 2014) were used, assuming the common ancestor of the order Capnodiales to be constrained to a minimum age of 100 MYA (gamma distribution, offset 100, mean 180, maximum soft bound 400). Diversification ages estimated using mitochondrial markers and bayesian analysis using both relaxed and strict clock models were younger than those calculated by Chang et al. (2016) using nuclear markers, penalized maximum likelihood analysis and strict clock, except for *Sphaerullina*. Divergence times were also estimated using strict clock to validate differences between mean node ages using relaxed lognormal clock versus strict clock.

For a strict clock, 95% highest posterior density (HPD) intervals were significantly broader (27-131 MY) in comparison to (6-23 MY) and mean node ages were among 9 – 43 MY older (Appendix A16, Table 4). Giving the low taxonomic sampling of this study is possible that diversification times can change when adding more taxa to the phylogeny. However, we estimate this change will not be dramatic.

**Tabla 4.**
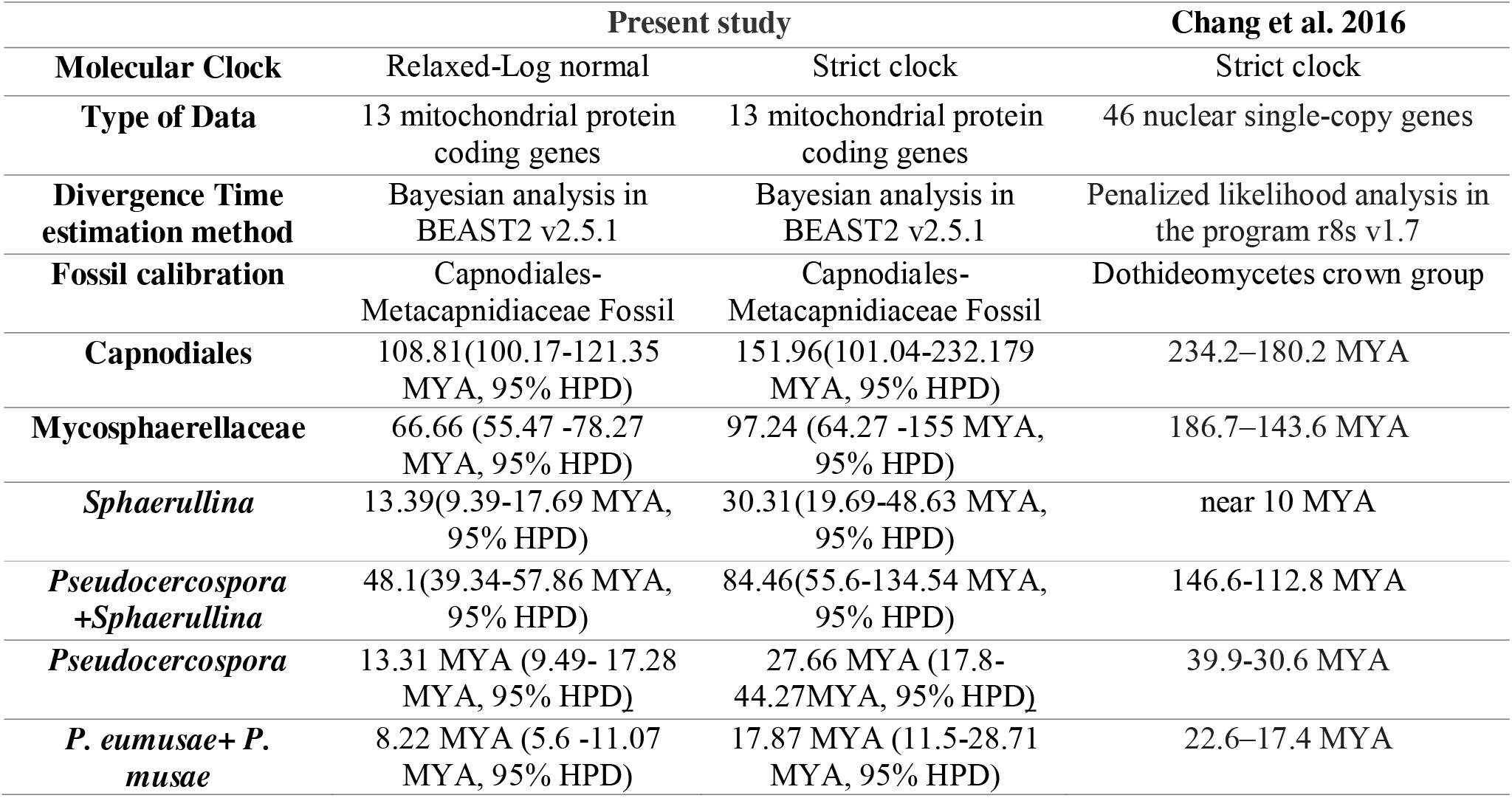
Comparison among Divergence time of some clades among different studies.

|  | **Present study** |  | **Chang et al. 2016** |
| --- | --- | --- | --- |
| **Molecular Clock** | Relaxed-Log normal | Strict clock | Strict clock |
| **Type of Data** | 13 mitochondrial protein coding genes | 13 mitochondrial protein coding genes | 46 nuclear single-copy genes |
| **Divergence Time estimation method** | Bayesian analysis in BEAST2 v2.5.1 | Bayesian analysis in BEAST2 v2.5.1 | Penalized likelihood analysis in the program r8s v1.7 |
| **Fossil calibration** | Capnodiales-Metacapnidiaceae Fossil | Capnodiales-Metacapnidiaceae Fossil | Dothideomycetes crown group |
| **Capnodiales** | 108.81(100.17-121.35 MYA, 95% HPD) | 151.96(101.04-232.179 MYA, 95% HPD) | 234.2–180.2 MYA |
| **Mycosphaerellaceae** | 66.66 (55.47 -78.27 MYA, 95% HPD) | 97.24 (64.27 -155 MYA, 95% HPD) | 186.7–143.6 MYA |
| ***Sphaerullina*** | 13.39(9.39-17.69 MYA, 95% HPD) | 30.31(19.69-48.63 MYA, 95% HPD) | near 10 MYA |
| ***Pseudocercospora* +*Sphaerullina*** | 48.1(39.34-57.86 MYA, 95% HPD) | 84.46(55.6-134.54 MYA, 95% HPD) | 146.6-112.8 MYA |
| ***Pseudocercospora*** | 13.31 MYA (9.49- 17.28 MYA, 95% HPD) | 27.66 MYA (17.8-44.27MYA, 95% HPD) | 39.9-30.6 MYA |
| ***P. eumusae*+ *P. musae*** | 8.22 MYA (5.6 -11.07 MYA, 95% HPD) | 17.87 MYA (11.5-28.71 MYA, 95% HPD) | 22.6–17.4 MYA |

## 5. Conclusions and Future directions

Mitochondrial genomes in Mycosphaerellaceae are highly variable in size and gene order, due to at least five independent intron invasion and horizontal gene transfer during their evolutionary history. Presence of duplicated genes, homing endonuclease genes (HEG) invasion and accessory genes was observed. Despite their order variability some genes were always recovered as neighbors in all mt genomes analyzed showing polycistronic transcription due to their endosymbiotic origin from an α-proteobacteria. Further gene editing and virulence assays will be important to shed light on fungal adaptation and more effective disease control strategies. Our mt phylogeny did not differ to previously published nuclear topologies. Fossil calibrated phylogenies reported for first time here for fungal plant pathogens had earlier diversification times for the origin of all the species involved compared to previous studies. Phylogenomic studies including a good taxonomic sampling and biogeographical analysis for *Pseudocercospora* species will further clarify whether the Sigatoka disease causing species virulence flare up after *Musa* domestication.

## Abbreviations

Mt genome: Mitochondrial genome
ETC genes: Electron transport chain genes
ORFs: Open reading frames
rns: Small ribosomal subunit RNA
rnl: Large ribosomal subunit rRNA
tRNAs: Transfer RNAs
HEG: Homing endonuclease genes
CDS: Coding DNA Sequence
HPD: Highest posterior density
MYA: Million years ago

## Data Availability Statement

The annotated mitochondrial genome of *Pseudocercospora fijiensis*, *atp6-atp8* and *nad2-nad3* sequences will be accessible at GeneBank upon paper acceptance. Phylogeny xml archive and other mitochondrial annotated genomes are available at https://sites.google.com/site/tatianatatianaarias/publications/arcila-et-al-supplementary-materials

## Acknowledgements

We thank Michael R. McKain, Yesid Cuesta Atroz and Diego Mauricio Riaño Pachón for their valuable advice during the course of the study, Luis Eduardo Mejia for help with illustrations, Alejandro Rodriguez Cabal for his guidance in Linux and bioinformatics tools, Isabel Cristina Calle for experimental assistance doing PCR assays and Juan Santiago Zuluaga for financial help to support Juliana Arcila work at CIB. Sequence data for *Pseudovirgaria hyperparasitica* were produced by the US Department of Energy Joint Genome Institute http://www.jgi.doe.gov/ in collaboration with the user community.

## Appendix A

Supplementary material data associated with this article can be found, in the online version, at XXXXXXX

## Bibiography

Adams, K. L. and Palmer, J. D. (2003) ‘Evolution of mitochondrial gene content: gene loss and transfer to the nucleus’, Molecular Phylogenetics and Evolution, 29(3), pp. 380–395. doi: 10.1016/S1055-7903(03)00194-5.

Aguileta, G. et al. (2014) ‘High variability of mitochondrial gene order among fungi’, Genome Biology and Evolution. doi: 10.1093/gbe/evu028.

Alverson, A. J. et al. (2010) ‘Insights into the Evolution of Mitochondrial Genome Size from Complete Sequences of Citrullus lanatus and Cucurbita pepo (Cucurbitaceae)’, Molecular Biology and Evolution. Oxford University Press, 27(6), pp. 1436–1448. doi: 10.1093/molbev/msq029.

Andrews S. (2010) ‘FastQC: A quality control tool for high throughput sequence data. Reference Source.’

Arango Isaza, R. E. et al. (2016) ‘Combating a Global Threat to a Clonal Crop: Banana Black Sigatoka Pathogen Pseudocercospora fijiensis (Synonym Mycosphaerella fijiensis) Genomes Reveal Clues for Disease Control’, PLOS Genetics. Edited by J. M. McDowell. Wiley-VCH Verlag GmbH & Co. KGaA, 12(8), p. e1005876. doi: 10.1371/journal.pgen.1005876.

Arzanlou, M., Groenewald, J. Z., Fullerton, R. A., Abeln, E. C. A., Carlier, J., Zapater, M.-F., et al. (2008) ‘Multiple gene genealogies and phenotypic characters differentiate several novel species of Mycosphaerella and related anamorphs on banana’, Persoonia, 20, pp. 19–37. doi: 10.3767/003158508X302212.

Arzanlou, M., Groenewald, J. Z., Fullerton, R. A., Abeln, E. C. A., Carlier, J., Zapater, M., et al. (2008) ‘Multiple gene genealogies and phenotypic characters differentiate several novel species of Mycosphaerella and related anamorphs on banana’, pp. 19–37. doi: 10.3767/003158508X302212.

Bankevich, A. et al. (2012) ‘SPAdes: A New Genome Assembly Algorithm and Its Applications to Single-Cell Sequencing’, Journal of Computational Biology, 19(5), pp. 455–477. doi: 10.1089/cmb.2012.0021.

Barbrook, A. C. et al. (2010) ‘Organization and expression of organellar genomes.’, Philosophical transactions of the Royal Society of London. Series B, Biological sciences. The Royal Society, 365(1541), pp. 785–97. doi: 10.1098/rstb.2009.0250.

Basse, C. W. (2010) ‘Mitochondrial inheritance in fungi’, Current Opinion in Microbiology, 13(6), pp. 712–719. doi: 10.1016/j.mib.2010.09.003.

Benson, D. A. et al. (1999) ‘GenBank’, Nucleic Acids Research. Oxford University Press, 27(1), pp. 12–17. doi: 10.1093/nar/27.1.12.

Bernt, M., Braband, A., et al. (2013) ‘Genetic aspects of mitochondrial genome evolution’, Molecular Phylogenetics and Evolution, 69(2), pp. 328–338. doi: 10.1016/j.ympev.2012.10.020.

Bernt, M., Donath, A., et al. (2013) ‘MITOS: Improved de novo metazoan mitochondrial genome annotation’, Molecular Phylogenetics and Evolution, 69(2), pp. 313–319. doi: 10.1016/j.ympev.2012.08.023.

Boetzer, M. et al. (2011) ‘Scaffolding pre-assembled contigs using SSPACE’, Bioinformatics. Oxford University Press, 27(4), pp. 578–579. doi: 10.1093/bioinformatics/btq683.

Bolger, A. M., Lohse, M. and Usadel, B. (2014) ‘Trimmomatic: a flexible trimmer for Illumina sequence data’, Bioinformatics. Oxford University Press, 30(15), pp. 2114–2120. doi: 10.1093/bioinformatics/btu170.

Bouckaert, R. et al. (2014) ‘BEAST 2: A Software Platform for Bayesian Evolutionary Analysis’, PLoS Computational Biology. Edited by A. Prlic. Public Library of Science, 10(4), p. e1003537. doi: 10.1371/journal.pcbi.1003537.

Braun, U. et al. (2011) ‘Pseudovirgaria, a fungicolous hyphomycete genus.’, IMA fungus. CBS Fungal Biodiversity Centre, 2(1), pp. 65–9. doi: 10.5598/imafungus.2011.02.01.09.

Burger, G., Gray, M. W. and Lang, B. F. (2003) ‘Mitochondrial genomes: Anything goes’, Trends in Genetics. doi: 10.1016/j.tig.2003.10.012.

Camacho, C. et al. (2009) ‘BLAST+: architecture and applications’, BMC Bioinformatics. BioMed Central, 10(1), p. 421. doi: 10.1186/1471-2105-10-421.

Cañas-Gutiérrez, G. P. et al. (2006) ‘Molecular characterization of benomyl-resistant isolates of Mycosphaerella fijiensis, collected in Colombia’, Journal of Phytopathology. doi: 10.1111/j.1439-0434.2006.01113.x.

Cañas-Gutiérrez, G. P. et al. (2009) ‘Analysis of the CYP51 gene and encoded protein in propiconazole-resistant isolates of *Mycosphaerella fijiensis*’, *Pest Management Science*. John Wiley & Sons, Ltd., 65(8), pp. 892–899. doi: 10.1002/ps.1770.

Carlier, J. et al. (2000) ‘Septoria Leaf Spot of Banana: A Newly Discovered Disease Caused by *Mycosphaerella eumusae* (Anamorph *Septoria eumusae*)’, Phytopathology. The American Phytopathological Society, 90(8), pp. 884–890. doi: 10.1094/PHYTO.2000.90.8.884.

Chang, T. C. et al. (2016) ‘Comparative Genomics of the Sigatoka Disease Complex on Banana Suggests a Link between Parallel Evolutionary Changes in Pseudocercospora fijiensis and Pseudocercospora eumusae and Increased Virulence on the Banana Host’, PLoS Genetics. doi: 10.1371/journal.pgen.1005904.

Chevalier, B. S. and Stoddard, B. L. (2001) ‘Homing endonucleases: structural and functional insight into the catalysts of intron/intein mobility.’, Nucleic acids research, 29(18), pp. 3757–74. doi: https://doi.org/10.1093/nar/29.18.3757.

Chrzanowska-Lightowlers, Z. (2015) ‘Mitochondrial RNA and its eccentricities’, Biochemist, 37(2), pp. 28–32. Available at: https://www.researchgate.net/publication/281799493_Mitochondrial_RNA_and_its_eccentricities (Accessed: 5 November 2018).

Churchill, A. C. L. (2011) ‘Mycosphaerella fijiensis, the black leaf streak pathogen of banana: Progress towards understanding pathogen biology and detection, disease development, and the challenges of control’, Molecular Plant Pathology. doi: 10.1111/j.1364-3703.2010.00672.x.

Clark-Walker, G. D. (1992) ‘Evolution of Mitochondrial Genomes in Fungi’, in, pp. 89–127. doi: 10.1016/S0074-7696(08)62064-1.

Conant, G. C. and Wolfe, K. H. (2008) ‘Turning a hobby into a job: How duplicated genes find new functions’, Nature Reviews Genetics. Nature Publishing Group, 9(12), pp. 938–950. doi: 10.1038/nrg2482.

Cools, H. J. and Fraaije, B. A. (2013) ‘Update on mechanisms of azole resistance in *Mycosphaerella graminicola* and implications for future control’, *Pest Management Science*. John Wiley & Sons, Ltd, 69(2), pp. 150–155. doi: 10.1002/ps.3348.

Darriba, D. et al. (2012) ‘jModelTest 2: more models, new heuristics and parallel computing’, Nature Methods. Nature Research, 9(8), pp. 772–772. doi: 10.1038/nmeth.2109.

Dassa, B. et al. (2009) ‘Fractured genes: A novel genomic arrangement involving new split inteins and a new homing endonuclease family’, Nucleic Acids Research. doi: 10.1093/nar/gkp095.

Depotter, J. et al. (2018) ‘Nuclear and mitochondrial genomes of the hybrid fungal plant pathogen Verticillium longisporum display a mosaic structure’, bioRxiv. Cold Spring Harbor Laboratory, p. 249565. doi: 10.1101/249565.

Dhillon, B. et al. (2015) ‘Horizontal gene transfer and gene dosage drives adaptation to wood colonization in a tree pathogen’, Proceedings of the National Academy of Sciences, 112(11), pp. 3451–3456. doi: 10.1073/pnas.1424293112.

Drenkhan, R. et al. (2014) ‘*Dothistroma septosporum* on firs (*Abies* spp.) in the northern Baltics’, Forest Pathology. Edited by J. Stenlid, 44(3), pp. 250–254. doi: 10.1111/efp.12110.

Drummond, A. J. et al. (2006) ‘Relaxed Phylogenetics and Dating with Confidence’. doi: 10.1371/journal.pbio.0040088.

Edgar, R. C. (2004) ‘MUSCLE: multiple sequence alignment with high accuracy and high throughput’, Nucleic Acids Research. Oxford University Press, 32(5), pp. 1792–1797. doi: 10.1093/nar/gkh340.

Finn, R. D. et al. (2014) ‘Pfam: the protein families database’, Nucleic Acids Research. Oxford University Press, 42(D1), pp. D222–D230. doi: 10.1093/nar/gkt1223.

Franz Lang, B., Laforest, M.-J. and Burger, G. (2007) ‘Mitochondrial introns: a critical view’. doi: 10.1016/j.tig.2007.01.006.

Giordano, L. et al. (2018) ‘Mitonuclear interactions may contribute to fitness of fungal hybrids’, Scientific Reports. Nature Publishing Group, 8(1), p. 1706. doi: 10.1038/s41598-018-19922-w.

Gonthier, P. et al. (2018) A fungal invasion is enhanced by hybridization and gene introgression: ecological and evolutionary implications of genomic admixing Presenting Author Co-authors Recording (mp4) Recording. Available at: https://iris.unito.it/retrieve/handle/2318/1695582/488831/GonthieretalICPP2018_invitedpresentation.pdf (Accessed: 29 April 2019).

Goodwin, S. B. et al. (2016) ‘The mitochondrial genome of the ethanol-metabolizing, wine cellar mold Zasmidium cellare is the smallest for a filamentous ascomycete’, Fungal Biology. doi: 10.1016/j.funbio.2016.05.003.

Grabherr, M. G. et al. (2011) ‘Full-length transcriptome assembly from RNA-Seq data without a reference genome’, Nature Biotechnology, 29(7), pp. 644–652. doi: 10.1038/nbt.1883.

Griffiths, A. J. (1995) ‘Natural plasmids of filamentous fungi.’, Microbiological reviews, 59(4), pp. 673–85. Available at: http://www.ncbi.nlm.nih.gov/pubmed/8531891 (Accessed: 28 November 2017).

Grigoriev, I. V. et al. (2014) ‘MycoCosm portal: gearing up for 1000 fungal genomes’, Nucleic Acids Research. Oxford University Press, 42(D1), pp. D699–D704. doi: 10.1093/nar/gkt1183.

Gualberto, J. M. et al. (2014) ‘The plant mitochondrial genome: Dynamics and maintenance’, Biochimie, 100, pp. 107–120. doi: 10.1016/j.biochi.2013.09.016.

Gueidan, C. et al. (2011) ‘Rock-inhabiting fungi originated during periods of dry climate in the late Devonian and middle Triassic.’, Fungal biology, 115(10), pp. 987–96. doi: 10.1016/j.funbio.2011.04.002.

Guindon, S. et al. (2010) ‘New Algorithms and Methods to Estimate Maximum-Likelihood Phylogenies: Assessing the Performance of PhyML 3.0’, *Systematic Biology*. The University of Texas at Austin, [Austin (TX)], 59(3), pp. 307–321. doi: 10.1093/sysbio/syq010.

Guindon, S., Gascuel, O. and Rannala, B. (2003) ‘A Simple, Fast, and Accurate Algorithm to Estimate Large Phylogenies by Maximum Likelihood’, Systematic Biology. Oxford University Press, 52(5), pp. 696–704. doi: 10.1080/10635150390235520.

Hane, J. K. et al. (2007) ‘Dothideomycete plant interactions illuminated by genome sequencing and EST analysis of the wheat pathogen Stagonospora nodorum.’, The Plant cell. American Society of Plant Biologists, 19(11), pp. 3347–68. doi: 10.1105/tpc.107.052829.

Hausner, G. (2003) ‘Fungal mitochondrial genomes, plasmids and introns’, Applied Mycology and Biotechnology. doi: 10.1016/S1874-5334(03)80009-6.

Hunt, M. et al. (2013) ‘REAPR: a universal tool for genome assembly evaluation’, Genome Biology. BioMed Central, 14(5), p. R47. doi: 10.1186/gb-2013-14-5-r47.

Jelen, V. et al. (2016) ‘Complete mitochondrial genome of the Verticillium-wilt causing plant pathogen Verticillium nonalfalfae’, PLOS ONE. Edited by M. Nowrousian. Public Library of Science, 11(2), p. e0148525. doi: 10.1371/journal.pone.0148525.

Joardar, V. et al. (2012) ‘Sequencing of mitochondrial genomes of nine Aspergillus and Penicillium species identifies mobile introns and accessory genes as main sources of genome size variability’, BMC Genomics, 13(1), p. 698. doi: 10.1186/1471-2164-13-698.

Johnson, M. et al. (2008) ‘NCBI BLAST: a better web interface’, Nucleic Acids Research, 36(Web Server), pp. W5–W9. doi: 10.1093/nar/gkn201.

Kawano, S., Takano, H. and Kuroiwa, T. (1995) ‘Sexuality of Mitochondria: Fusion, Recombination, and Plasmids’, in, pp. 49–110. doi: 10.1016/S0074-7696(08)62496-1.

Kearse, M. et al. (2012) ‘Geneious Basic: An integrated and extendable desktop software platform for the organization and analysis of sequence data’, Bioinformatics. Oxford University Press, 28(12), pp. 1647–1649. doi: 10.1093/bioinformatics/bts199.

Kitchen, J. L. et al. (2016) ‘The Evolution of Fungicide Resistance Resulting from Combinations of Foliar-Acting Systemic Seed Treatments and Foliar-Applied Fungicides: A Modeling Analysis’, PLOS ONE. Edited by P. Heneberg. Clarendon Press, 11(8), p. e0161887. doi: 10.1371/journal.pone.0161887.

Ladoukakis, E. D. and Zouros, E. (2017) ‘Evolution and inheritance of animal mitochondrial DNA: rules and exceptions’, Ladoukakis and Zouros J of Biol Res-Thessaloniki, 24(2). doi: 10.1186/s40709-017-0060-4.

De Lapeyre, L. et al. (2010) ‘Black Leaf Streak Disease is challenging the banana industry’, XXXX, 65(65), pp. 327–342. doi: 10.1051/fruits/2010034.

Larkin, M. A. et al. (2007) ‘Clustal W and Clustal X version 2.0’, Bioinformatics. Oxford University Press, 23(21), pp. 2947–2948. doi: 10.1093/bioinformatics/btm404.

Lavrov, D. V. and Pett, W. (2016) ‘Animal Mitochondrial DNA as We Do Not Know It: mt-Genome Organization and Evolution in Nonbilaterian Lineages’, Genome Biology and Evolution, 8(9), pp. 2896–2913. doi: 10.1093/gbe/evw195.

Lodish, H. F. (2003) Molecular cell biology. 5th edn. W.H. Freeman. Available at: https://www.ncbi.nlm.nih.gov/books/NBK21475/ (Accessed: 24 May 2017).

Ma, Z. and Michailides, T. J. (2005) ‘Advances in understanding molecular mechanisms of fungicide resistance and molecular detection of resistant genotypes in phytopathogenic fungi’, Crop Protection, 24(10), pp. 853–863. doi: 10.1016/j.cropro.2005.01.011.

Mardanov, A. V et al. (2014) ‘The 203 kbp mitochondrial genome of the phytopathogenic fungus Sclerotinia borealis reveals multiple invasions of introns and genomic duplications.’, PloS one. Public Library of Science, 9(9), p. e107536. doi: 10.1371/journal.pone.0107536.

Mi, H. et al. (2017) ‘PANTHER version 11: expanded annotation data from Gene Ontology and Reactome pathways, and data analysis tool enhancements’, Nucleic Acids Research. Oxford University Press, 45(D1), pp. D183–D189. doi: 10.1093/nar/gkw1138.

Nadalin, F., Vezzi, F. and Policriti, A. (2012) ‘GapFiller: a de novo assembly approach to fill the gap within paired reads’, BMC Bioinformatics. BioMed Central, 13(Suppl 14), p. S8. doi: 10.1186/1471-2105-13-S14-S8.

O’Leary, N. A. et al. (2016) ‘Reference sequence (RefSeq) database at NCBI: current status, taxonomic expansion, and functional annotation’, Nucleic Acids Research, 44(D1), pp. D733–D745. doi: 10.1093/nar/gkv1189.

Ohm, R. A. et al. (2012) ‘Diverse Lifestyles and Strategies of Plant Pathogenesis Encoded in the Genomes of Eighteen Dothideomycetes Fungi’, PLoS Pathogens. doi: 10.1371/journal.ppat.1003037.

Okorski, A. et al. (2016) ‘The complete mitogenome of *Mycosphaerella pinodes* (Ascomycota, Mycosphaerellaceae)’, Mitochondrial DNA Part B. Taylor & Francis, 1(1), pp. 48–49. doi: 10.1080/23802359.2015.1137817.

Paquin, B. et al. (1997) ‘The fungal mitochondrial genome project: evolution of fungal mitochondrial genomes and their gene expression.’, Current genetics, 31(5), pp. 380–95. doi: https://doi.org/10.1007/s002940050220.

Rambaut, A. (2006) ‘FigTree. Tree figure drawing tool version 1.3. 1’. Institute of Evolutionary biology, University of Edinburgh.

Rambaut, A. and Drummond, A. (2009) ‘Tracer: MCMC trace analysis tool version 1.5’. Oxford: University of Oxford.

Rosewich, U. L. and Kistler, H. C. (2000) ‘Role of Horizontal Gene Transfer in the Evolution of Fungi’, Annual Review of Phytopathology, 38(1), pp. 325–363. doi: 10.1146/annurev.phyto.38.1.325.

Salavirta, H. et al. (2014) ‘Mitochondrial Genome of Phlebia radiata Is the Second Largest (156 kbp) among Fungi and Features Signs of Genome Flexibility and Recent Recombination Events’, PLoS ONE. Edited by J.-H. Yu. Public Library of Science, 9(5), p. e97141. doi: 10.1371/journal.pone.0097141.

Sanderson, M. J. (2003) ‘r8s: inferring absolute rates of molecular evolution and divergence times in the absence of a molecular clock.’, Bioinformatics (Oxford, England), 19(2), pp. 301–302. doi: https://doi.org/10.1093/bioinformatics/19.2.301.

Schäfer, B., Hansen, M. and Lang, B. F. (2005) ‘Transcription and RNA-processing in fission yeast mitochondria.’, RNA (New York, N.Y.). Cold Spring Harbor Laboratory Press, 11(5), pp. 785–95. doi: 10.1261/rna.7252205.

Schmidt, A. R. et al. (2014) ‘Amber fossils of sooty moulds’, Review of Palaeobotany and Palynology, 200, pp. 53–64. doi: 10.1016/j.revpalbo.2013.07.002.

Schwarz, G. (1978) ‘Estimating the Dimension of a Model’, The Annals of Statistics. Institute of Mathematical Statistics, 6(2), pp. 461–464. doi: 10.1214/aos/1176344136.

Shen, X.-Y. et al. (2015) ‘Characterization and Phylogenetic Analysis of the Mitochondrial Genome of Shiraia bambusicola Reveals Special Features in the Order of Pleosporales’, PLOS ONE. Edited by M.-J. Virolle. CAB International, 10(3), p. e0116466. doi: 10.1371/journal.pone.0116466.

Sierotzki, H. et al. (2000) ‘Mode of resistance to respiration inhibitors at the cytochrome bc1 enzyme complex ofMycosphaerella fijiensis field isolates’, *Pest Management Science*. John Wiley & Sons, Ltd., 56(10), pp. 833–841. doi: 10.1002/1526-4998(200010)56:10<833::AID-PS200>3.0.CO;2-Q.

Stamatakis, A. (2014) ‘RAxML version 8: a tool for phylogenetic analysis and post-analysis of large phylogenies’, Bioinformatics. Oxford University Press, 30(9), pp. 1312–1313. doi: 10.1093/bioinformatics/btu033.

Stukenbrock, E. H. et al. (2006) ‘Origin and Domestication of the Fungal Wheat Pathogen Mycosphaerella graminicola via Sympatric Speciation’, Molecular Biology and Evolution. Oxford University Press, Oxford, 24(2), pp. 398–411. doi: 10.1093/molbev/msl169.

Stukenbrock, E. H. and Mcdonald, B. A. (2008) ‘The Origins of Plant Pathogens in Agro-Ecosystems’, pp. 75–100. doi: 10.1146/annurev.phyto.010708.154114.

Thomma, B. P. H. J. et al. (2005) ‘Cladosporium fulvum (syn. Passalora fulva), a highly specialized plant pathogen as a model for functional studies on plant pathogenic Mycosphaerellaceae’, Molecular Plant Pathology. Blackwell Science Ltd, 6(4), pp. 379–393. doi: 10.1111/j.1364-3703.2005.00292.x.

Torriani, S. F. F. et al. (2008) ‘Intraspecific comparison and annotation of two complete mitochondrial genome sequences from the plant pathogenic fungus Mycosphaerella graminicola’, Fungal Genetics and Biology, 45(5), pp. 628–637. doi: 10.1016/j.fgb.2007.12.005.

Torriani, S. F. F. et al. (2014) ‘Comparative analysis of mitochondrial genomes from closely related Rhynchosporium species reveals extensive intron invasion’, Fungal Genetics and Biology. Academic Press, 62, pp. 34–42. doi: 10.1016/J.FGB.2013.11.001.

Videira, S. I. R. et al. (2017) ‘Mycosphaerellaceae – chaos or clarity?’, Studies in Mycology. doi: 10.1016/.

Wijayawardene, N. N. et al. (2014) ‘Naming and outline of Dothideomycetes-2014 including proposals for the protection or suppression of generic names.’, Fungal diversity. Europe PMC Funders, 69(1), pp. 1–55. doi: 10.1007/s13225-014-0309-2.

Wolf, K. and Giudice, L. Del (1988) ‘The Variable Mitochondrial Genome of Ascomycetes: Organization, Mutational Alterations, and Expression’, in Caspari, E. W. and Scandalios, J. G. (eds). Academic Press (Advances in Genetics), pp. 185–308. doi: https://doi.org/10.1016/S0065-2660(08)60460-5.

Zhang, S. (1995) The Structural Differences Between Animal and Plant Mitochondrial Genomes -- Two Evolutionary Scenarios, Zoological research. Available at: http://www.zoores.ac.cn/EN/Y1995/V16/I2/132 (Accessed: 28 April 2019).

