## Appendix for "Comparative genomics in plant fungal pathogens (Mycosphaerellaceae): variation in mitochondrial composition due to at least five independent intron invasions"

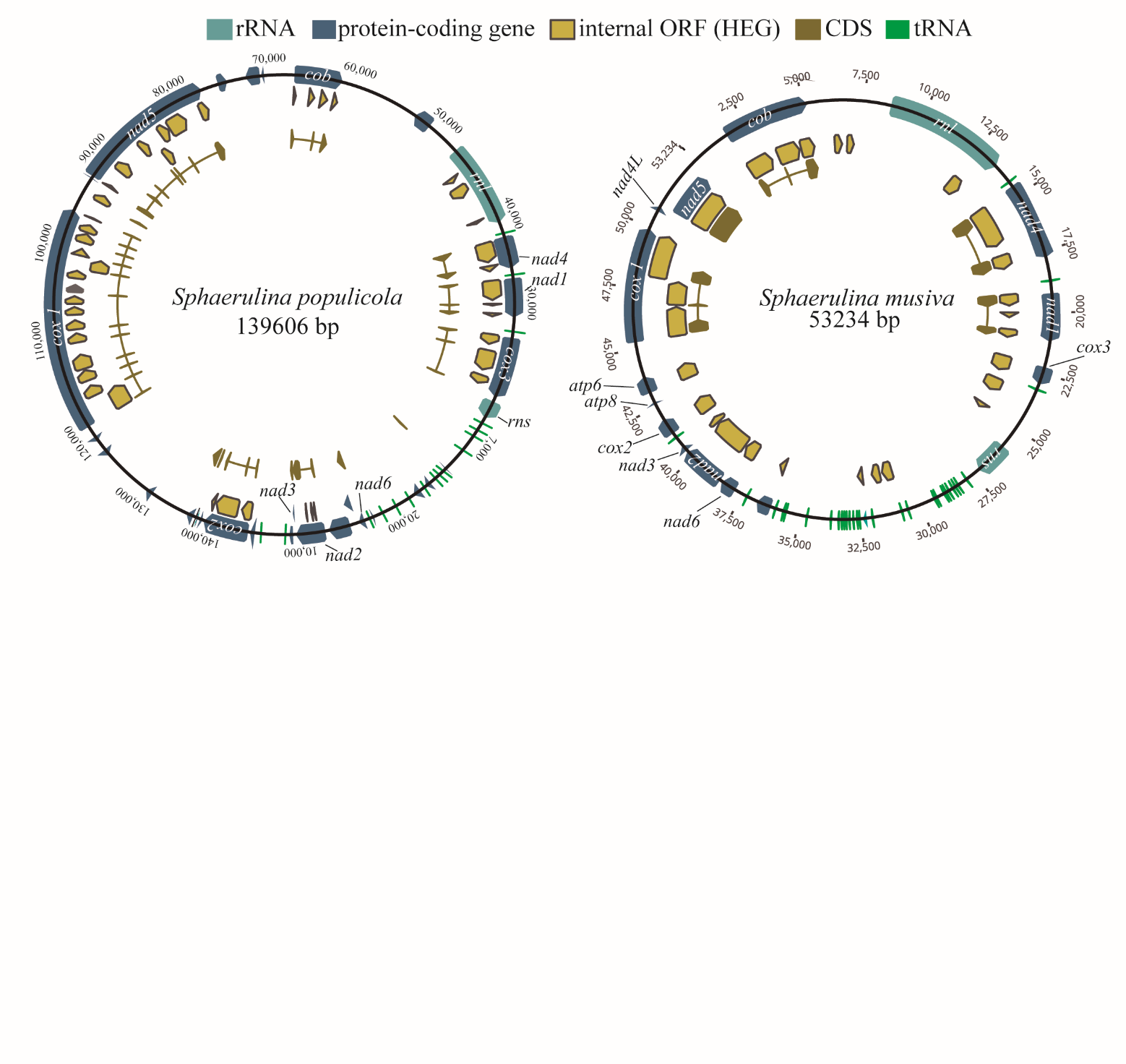

**Appendix 1. Genetic map of *S. musiva* and *S. populicola* mitochondrial genomes.** Two closely related species with the same core gene order but highly different size due to invasion of HEG related ORFS in *S. populicola*

**Appendix 2. Example of size increment in core genes due to introns.** Codifying size of *cob* gene of *P.fijiensis* is 130 pb but contains an intron that increases its size 1227 pb. A. Schematic representation of *cob* region in the genome. B. Intron presence confirmed by PCR.

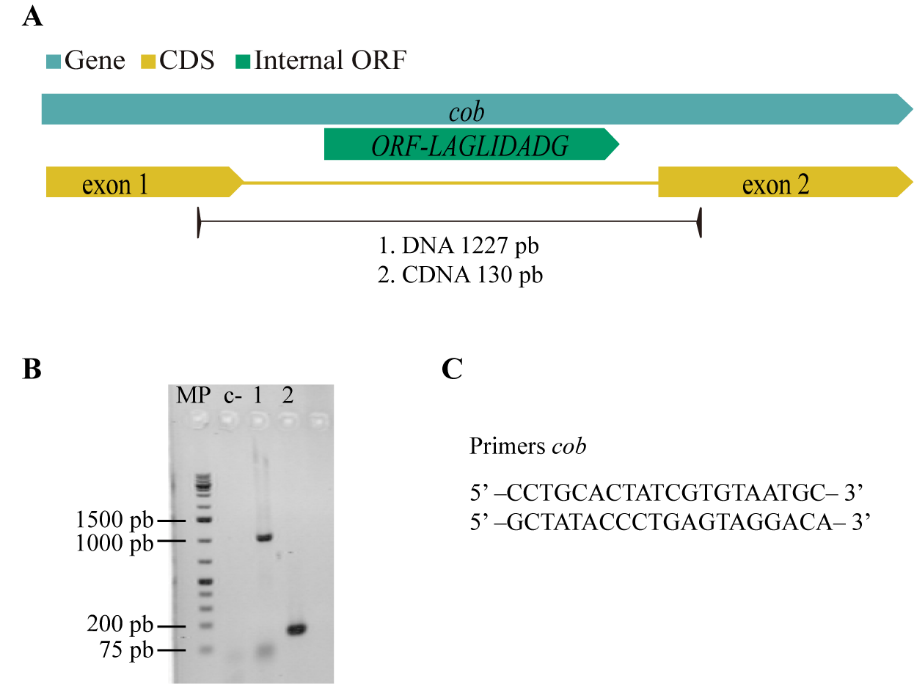

**Appendix 3. Maximum likelihood tree for *atp6*.**

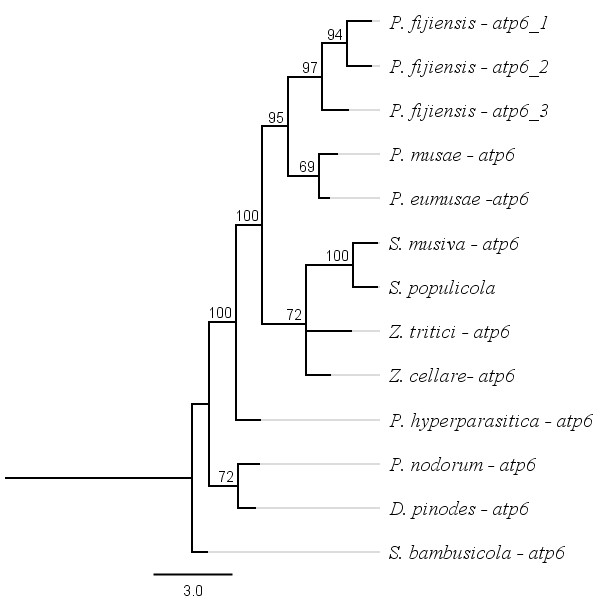

**Appendix 4. Maximum likelihood tree for *atp8*.**

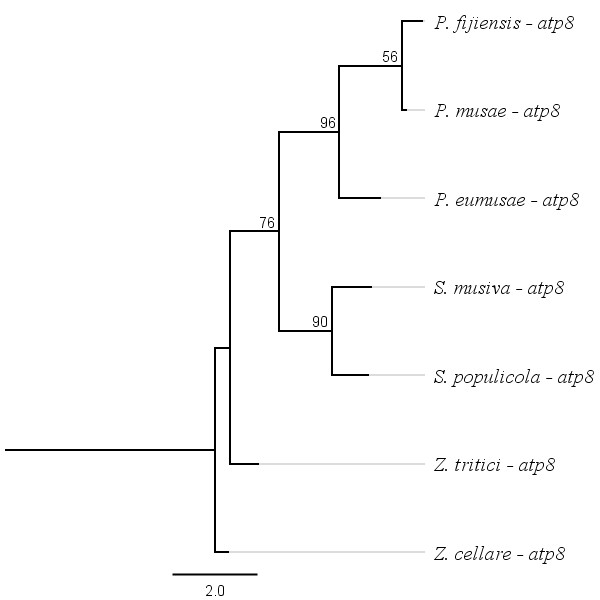

**Appendix 5. Maximum likelihood tree for *atp9*.**

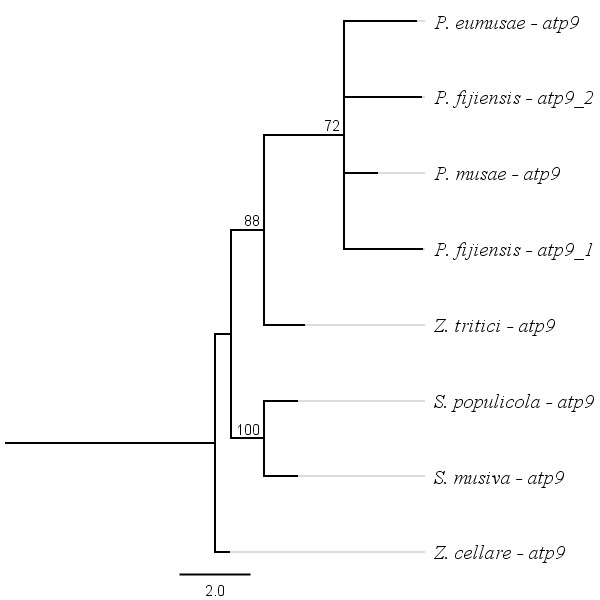

**Appendix 6. Maximum likelihood tree for *cox1*.**

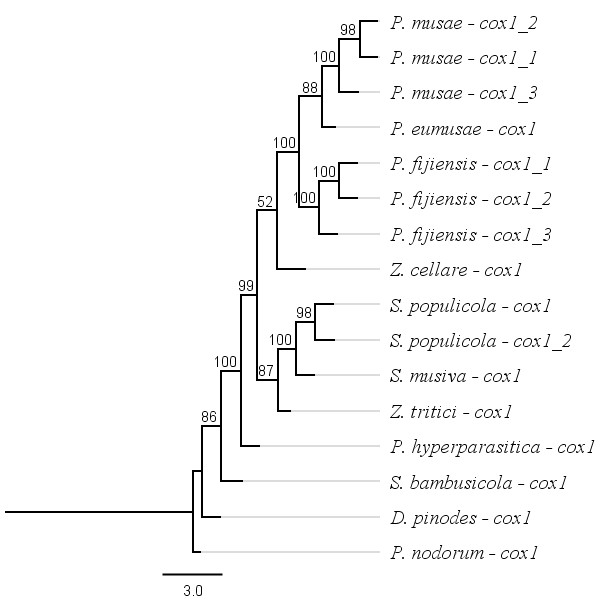

**Appendix 7. Maximum likelihood tree for *cox2*.**

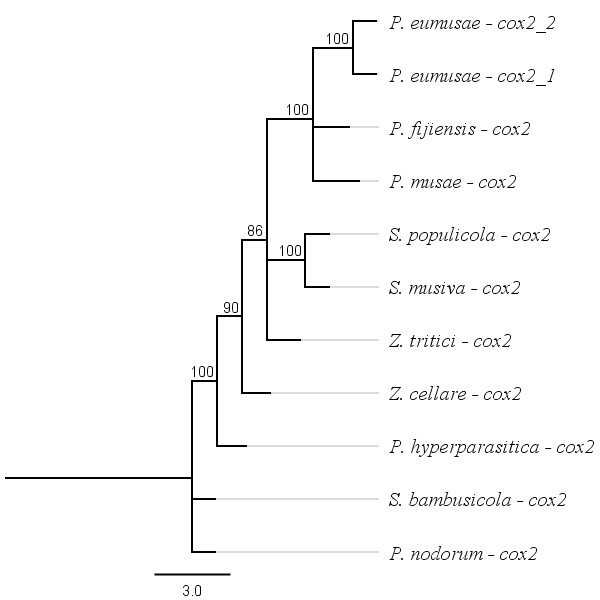

**Appendix 8. Maximum likelihood tree for *cox3.***

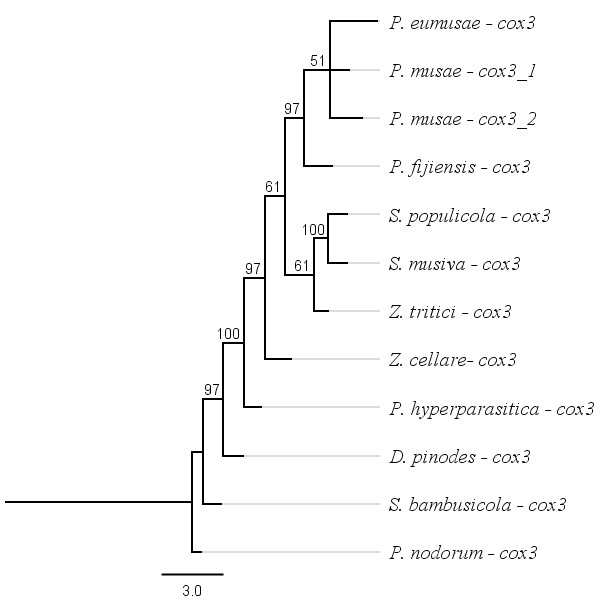

**Appendix 9. Maximum likelihood tree for *cob.***

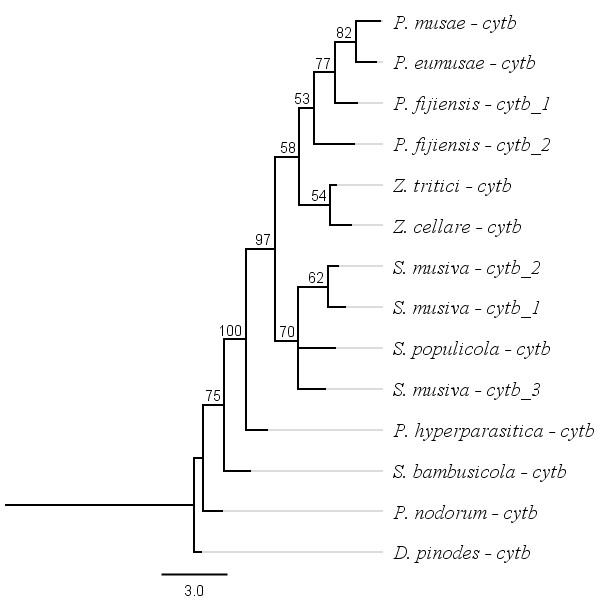

**Appendix 10. Maximum likelihood tree for *nad1.***

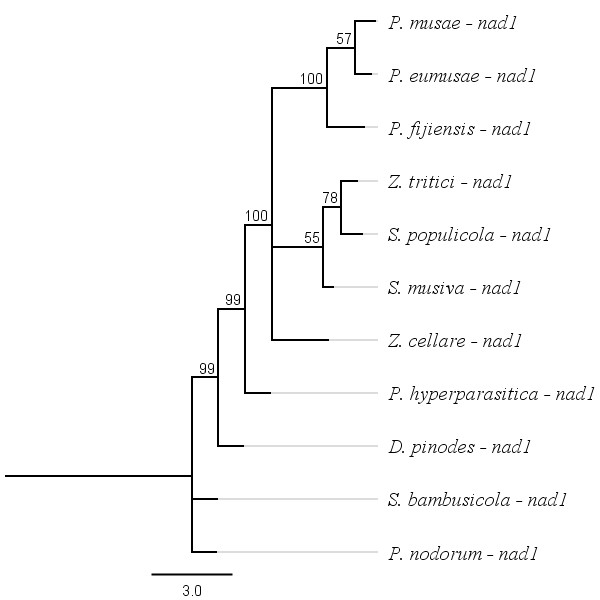

**Appendix 11. Maximum likelihood tree for *nad2.***

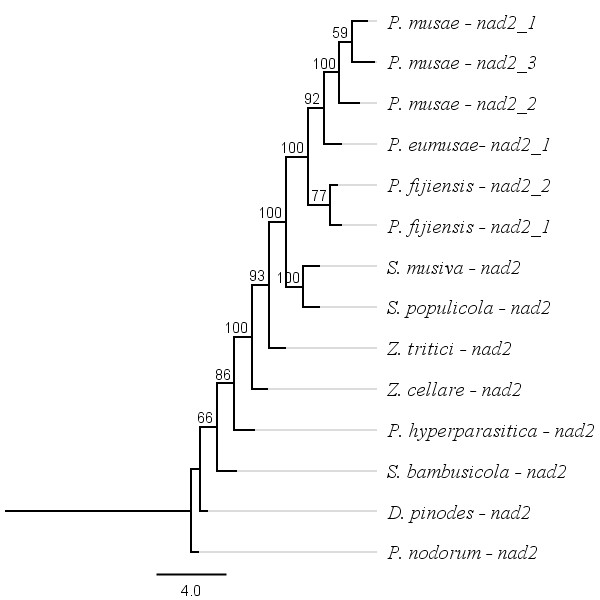

**Appendix 12. Maximum likelihood tree for *nad3.***

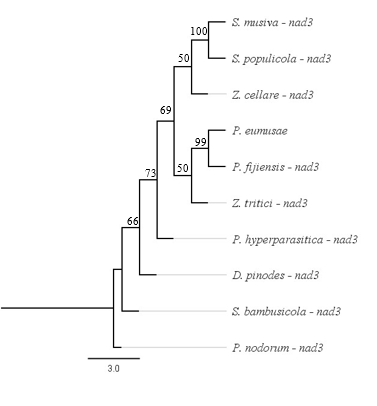

**Appendix 13. Maximum likelihood tree for *nad4.***

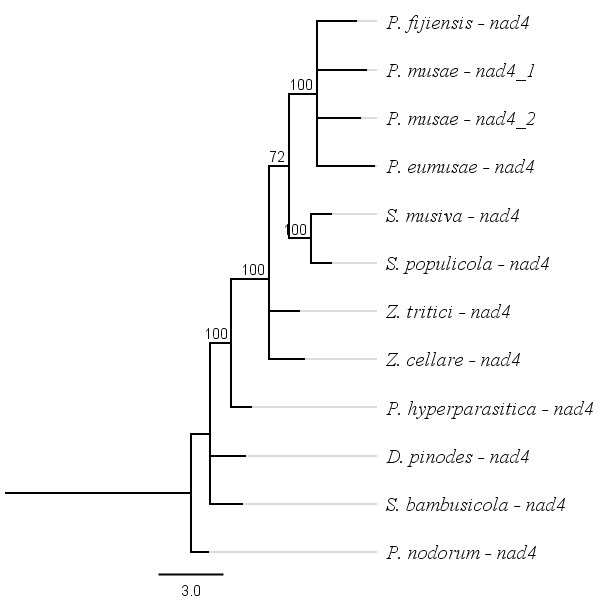

**Appendix 14. Maximum likelihood tree for *nad5.***

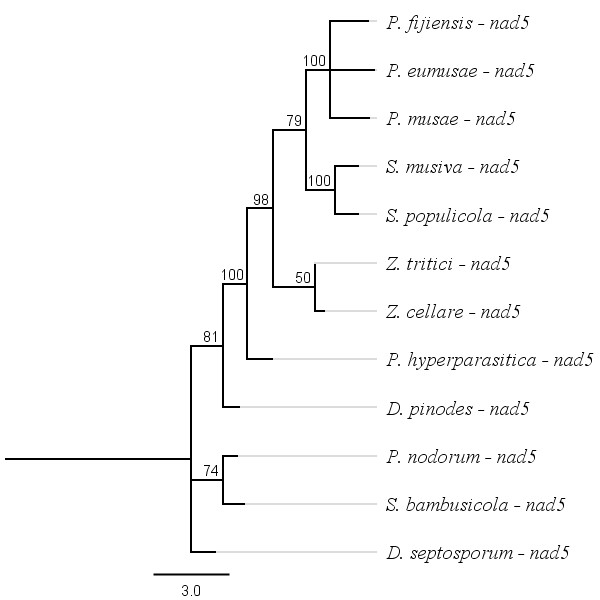

**Appendix 15. Maximum likelihood tree for *nad6.***

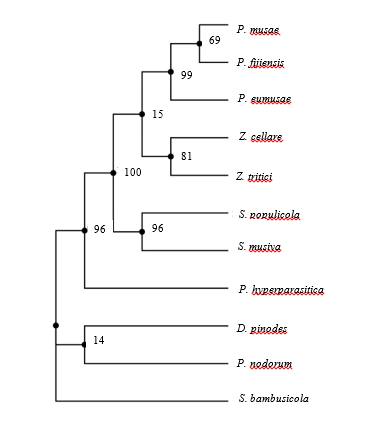

**Appendix 15. Maximum likelihood tree for *rns.***

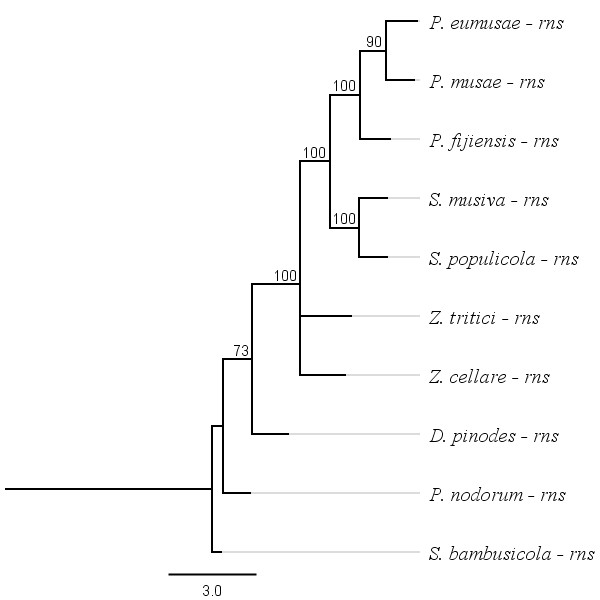

**Appendix 15. Bayesian phylogeny.** Using gamma prior for fossil calibration and strict clock

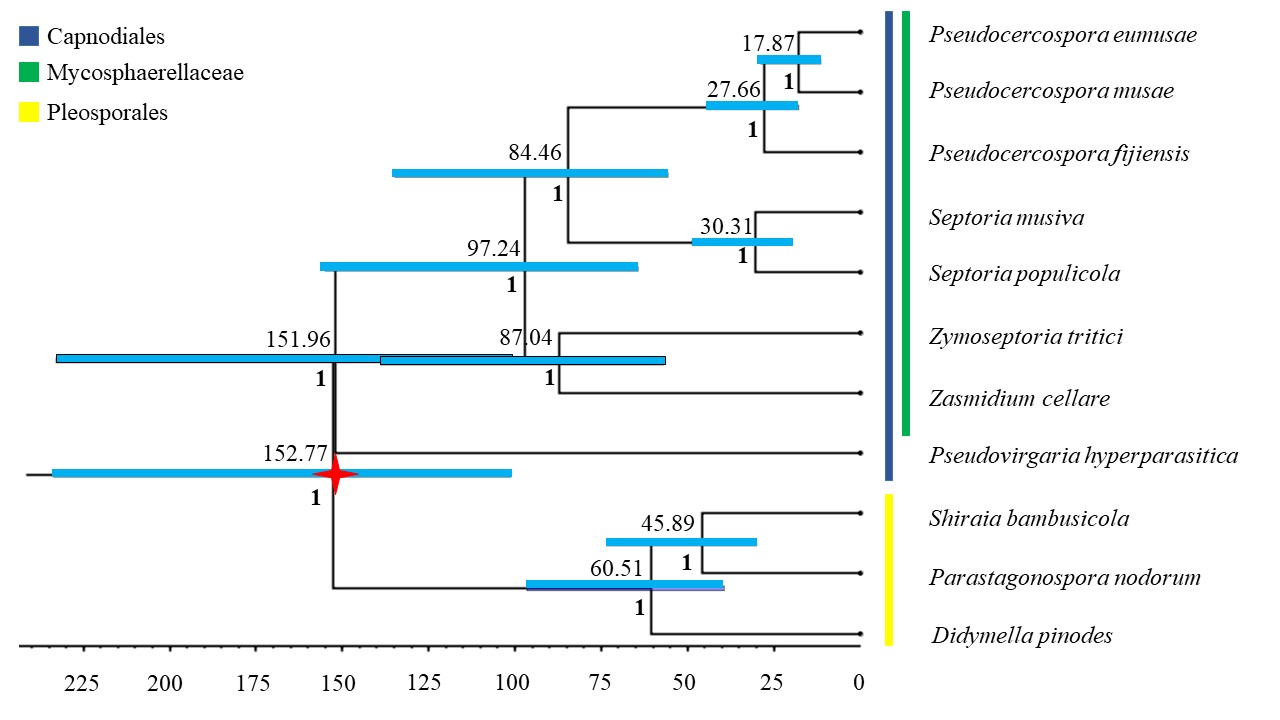
